# Mouse Model of Heart Attack and Stroke Shows Improved Survival with MPO Inhibition

**DOI:** 10.1101/2023.10.06.561304

**Authors:** Sohel Shamsuzzaman, Rebecca A. Deaton, Heather Doviak, Megan A. Evans, Anita Salamon, Santosh Karnewar, Vlad Serbulea, Gabriel F. Alencar, Laura S. Shankman, Kenneth Walsh, Stefan Bekiranov, Olivier Kocher, Monty Krieger, Bengt Kull, Marie Persson, Nils Bergenhem, Sepideh Heydarkhan-Hagvall, Gary K. Owens

**Author notes:** Corresponding author: Dr. Gary K. Owens, University of Virginia School of Medicine, Robert M. Berne Cardiovascular Research Center, PO Box 801394, Charlottesville, Virginia 22908-1394. These authors contributed equally to this work (Sohel Shamsuzzaman, Rebecca A. Deaton).

## Abstract

Thromboembolic events, including myocardial infarction (MI) or stroke, caused by the rupture or erosion of unstable atherosclerotic plaques are the leading cause of death worldwide^1^. Unfortunately, the lack of a mouse model that develops advanced coronary atherosclerosis and that exhibits a high incidence of spontaneous plaque rupture with MI or stroke has greatly stymied development of more effective therapeutic approaches for reducing these events beyond what has been achieved with aggressive lipid lowering. Herein, we describe a novel mouse model that develops widespread advanced atherosclerosis including in coronary, brachiocephalic, and carotid arteries. These mice show high mortality following Western Diet feeding with clear evidence of plaque rupture, MI, and stroke. To validate the utility of this model, mice were treated with the drug candidate AZM198, which inhibits myeloperoxidase, an enzyme primarily produced by activated neutrophils and predictive of rupture of human atherosclerotic lesions^2–7^. AZM198 treatment resulted in marked improvements in survival with a greater than 60% decrease in the incidence of plaque rupture, MI, and stroke. In summary, our work describes a novel mouse model that closely replicates late-stage clinical events of advanced human atherosclerotic disease and evidence that this model can be used to identify and test potential new therapeutic agents to prevent major adverse cardiac events.

## Main

Thromboembolic events caused by the rupture or erosion of unstable atherosclerotic plaques are the most common causes of cardiovascular morbidity and mortality^8,9^. Despite widespread use of statins, other lipid-lowering therapies, and lifestyle modifications, atherosclerosis-associated complications, including MI or stroke, remain the leading causes of death in the United States and worldwide^1^.

The most commonly used mouse models of atherosclerosis, such as *Apoe* and *Ldlr* knockout (KO) mice, develop plaques in the aortic root, aorta, and carotid arteries. However, they do not readily develop advanced coronary artery lesions even after extensive high-fat diet feeding^10–12^. They also do not undergo spontaneous plaque rupture with MI or stroke or do so at such a low frequency that they are not viable experimental models to study late-stage thrombotic events or to identify novel therapeutic approaches for treating the disease^10–12^. There are several reported mouse models of MI and plaque rupture^13–16,2,17–20^. However, none have been widely adopted or accepted as models of advanced human disease and/or they exhibit many unwanted side effects. A detailed comparison of many of these models was reported by Noonan *et al*^21^.

Perhaps the most interesting models involved the KO or mutation^17–20,22^ of the high density lipoprotein (HDL) receptor, SR-BI (encoded by *Scarb1*), that is involved in reverse cholesterol transport^23–25^. Genome-wide association studies (GWAS) have linked the human *SCARB1* locus with coronary artery disease (CAD) risk^26,27^. Moreover, recent human genetic studies have reported the Mendelian inheritance of CAD as a result of human *SCARB1* variants^28,29^. Consistent with these observations, diet-inducible coronary atherosclerosis was reported in SR-BI/*Ldlr* double KO mice and SR-BI KO/ApoeR61 mice with ≤12 weeks feeding with a variety of different atherogenic diets^18,30^. However, there are several major limitations to these models including female infertility and partially penetrant embryonic lethality in SR-BI KO mice. Pal *et al*. characterized a mouse model with a C-terminal three amino acid truncation mutant of SR-BI (SR-BI^ΔCT/ΔCT^) which interferes with the receptor’s binding to the adaptor protein PDZK1, dramatically reducing hepatic SR-BI expression and altering lipoprotein metabolism^19^. Unlike SR-BI KO mice, SR-BI^ΔCT/ΔCT^ mice are fertile^19^. However, when crossed to *Apoe*^−/−^ they suffer from rapid onset of disease with a median age of death at ∼8 weeks that is unlikely to model many aspects of late-stage human atherosclerotic coronary artery disease^19^.

Herein, we describe a novel mouse model that develops diet-inducible advanced atherosclerosis within coronary, brachiocephalic (BCA), and carotid arteries and which shows clear evidence of spontaneous plaque rupture, myocardial infarction, and stroke. In addition, we incorporated *Myh11-CreER^T^*^2^-eYFP transgenes into this mouse to permit smooth muscle cell (SMC) lineage tracing. The latter is important to enable assessment of SMC phenotypic transitions which we^31,32^ and others^33^ have shown play a critical role in late-stage lesion pathogenesis. The full genotype of the mouse is *Myh11-CreER^T^*^2^-eYFP SR-BI^ΔCT/ΔCT^ *Ldlr^−/−^* (henceforth abbreviated as “SR-BI^ΔCT/ΔCT^/*Ldlr^−/−^*”).

### High mortality of SR-BI^ΔCT/ΔCT^/*Ldlr^−/−^* mice fed Western Diet

SR-BI^ΔCT/ΔCT^/*Ldlr^−/−^* mice and their wild-type (SR-BI^WT/WT^/*Ldlr^−/−^*) and heterozygous (SR-BI^WT/ΔCT^/*Ldlr^−/−^*) littermate controls were fed a tamoxifen diet between 6-8 weeks of age to induce lineage tracing of SMCs. After tamoxifen feeding, mice were fed either a Chow or Western Diet (WD) during which time they were closely monitored for signs of decline, including hunched posture, labored breathing, hind-limb paralysis, and head tilt. As per our Animal Care and Use Committee (ACUC) approved protocol, mice exhibiting these symptoms were humanely euthanized and tissues were collected for analysis. A schematic of the experimental design is shown in **Figure 1a**. All the SR-BI^ΔCT/ΔCT^/*Ldlr^−/−^* mice and their wild-type (SR-BI^WT/WT^/*Ldlr^−/−^*) and heterozygous (SR-BI^WT/ΔCT^/*Ldlr^−/−^*) littermate controls showed 100% survival following 26 weeks of standard rodent Chow diet feeding (**Figure 1b-d**) and showed no signs of MI, stroke, or plaque rupture (**Extended Data Figure 1**). Chow-fed SR-BI^ΔCT/ΔCT^/*Ldlr^−/−^*mice had a 58% increase in baseline total cholesterol but no differences in triglycerides (TG) and displayed no significant differences in the liver enzymes alanine transaminase (ALT) and aspartate transaminase (AST) compared to SR-BI^WT/WT^/*Ldlr^−/−^* controls (**Extended Data Figure 2a**). There were also no significant differences in body weight, relative organ weight or total organ weight measurements compared to SR-BI^WT/WT^/*Ldlr^−/−^* (**Extended Data Figure 2b,c**).

**Figure 1:**
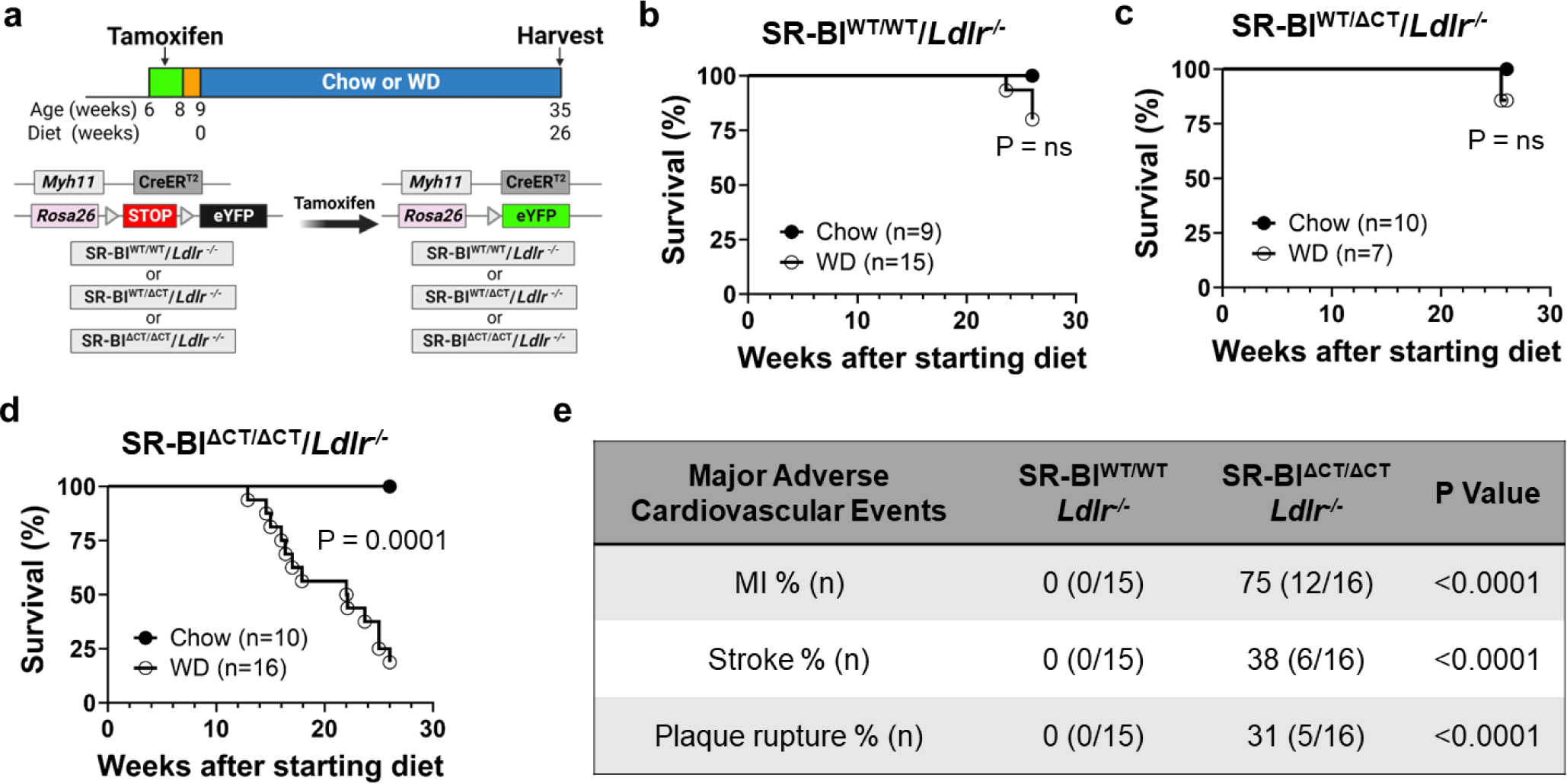
SR-BI^ΔCT/ΔCT^/*Ldlr^−/−^* mice fed a Western Diet show high mortality and increased Major Adverse Cardiovascular Events compared to SR-BI^WT/WT/^*Ldlr^−/−^* or SR-BI^WT/ΔCT^/*Ldlr^−/−^* mice. **(a)** Experimental design. Transient tamoxifen treatment induced expression of eYFP, a fluorescence marker for smooth muscle cell (SMC) lineage tracing. Mice were then fed either standard Chow or an atherogenic Western Diet (WD) for 26 weeks. Male mice were used in all experiments as the *Myh11-CreER^T^*^2^ transgene is located on the Y chromosome. **(b-d)** Kaplan-Meier survival curves (defined by humane terminal endpoint criteria, see Methods) for **(b)** SR-BI^WT/WT^/*Ldlr^−/−^* mice (Chow, n = 9 or WD, n = 15), **(c)** SR-BI^WT/ΔCT^/*Ldlr^−/−^* (Chow, n = 10 or WD, n = 7) or **(d)** SR-BI^ΔCT/ΔCT^/*Ldlr^−/−^* mice (Chow, n = 10 or WD, n = 16). **(e)** Table showing percentage of Major Adverse Cardiovascular Events in WD-fed SR-BI^WT/WT^/*Ldlr^−/−^* (n = 15) and SR-BI^ΔCT/ΔCT^/*Ldlr^−/−^*(n = 16) mice. Statistical analyses were performed using Log-rank (Mantel-Cox) test (**b**-**d**) and chi-square (Fisher’s exact) test (**e**). ns, not significant (P > 0.05). Panel **a** generated using BioRender.com.

When fed a Western Diet (WD) for 26 weeks, SR-BI^ΔCT/ΔCT^/*Ldlr^−/−^* mice showed a marked increase in mortality (81%, 13/16 mice) compared to SR-BI^WT/WT^/*Ldlr^−/−^* (20%, 3/15 mice) or SR-BI^WT/ΔCT^ /*Ldlr^−/−^* (14%, 1/7 mice) littermate controls (**Figure 1b-d**). The median time of death for WD-fed SR-BI^ΔCT/ΔCT^/*Ldlr^−/−^* (when 50% of the mice had met the humane terminal endpoint) was ∼20 weeks after initiating WD feeding. WD-fed heterozygous SR-BI^WT/ΔCT^/*Ldlr^−/−^* mice displayed similar mortality to WD-fed SR-BI^WT/WT^/*Ldlr^−/−^*littermate controls, indicating that one functional copy of SR-BI was sufficient to prevent diet-induced death. Thus, we compared SR-BI^ΔCT/ΔCT^/*Ldlr^−/−^* to SR-BI^WT/WT^/*Ldlr^−/−^* mice for the remainder of these studies. Consistent with the mortality data, none of the WD-fed SR-BI^WT/WT^/*Ldlr^−/−^* mice displayed signs of MI, stroke, or plaque rupture, whereas 75% of the WD-fed SR-BI^ΔCT/ΔCT^/*Ldlr^−/−^* mice had MI, 38% had stroke, and 31% had plaque rupture (**Figure 1e**). While markedly elevated compared to Chow-fed mice, WD-fed SR-BI^ΔCT/ΔCT^/*Ldlr^−/−^* mice had similar plasma levels of total cholesterol, TG, and liver enzymes (AST and ALT) compared to WD-fed SR-BI^WT/WT^/*Ldlr^−/−^* littermate controls (**Extended Data Figure 3a**). In addition, WD-fed SR-BI^ΔCT/ΔCT^/*Ldlr^−/−^*mice had significantly reduced body weight and markedly decreased epididymal (epi) fat (whether measured as a ratio to body weight or as absolute weight) compared to the WD-fed SR-BI^WT/WT^/*Ldlr^−/−^* littermate controls (**Extended Data Figure 3b,c**). WD-fed SR-BI^ΔCT/ΔCT^/*Ldlr^−/−^* mice also displayed cardiac hypertrophy and splenomegaly whether organs were measured as a ratio normalized to body weight or as absolute weights (**Extended Data Figure 3b,c**).

### Major Adverse Cardiovascular Events in Western Diet-fed mice

Tissues collected from SR-BI^ΔCT/ΔCT^/*Ldlr^−/−^*and SR-BI^WT/WT^/*Ldlr^−/−^* mice were examined macroscopically and histologically to determine whether the mice displayed coronary artery atherosclerosis and evidence of Major Adverse Cardiovascular Events (MACE) (including MI, plaque rupture or stroke). None of the SR-BI^ΔCT/ΔCT^/*Ldlr^−/−^*or SR-BI^WT/WT^/*Ldlr^−/−^* mice fed a standard rodent Chow diet for 26 weeks developed advanced atherosclerotic plaques or cardiac fibrosis (**Extended Data Figure 1**). In contrast, when fed a WD both the SR-BI^ΔCT/ΔCT^/*Ldlr^−/−^* and SR-BI^WT/WT^/*Ldlr^−/−^*mice exhibited advanced atherosclerotic plaques in the aortic sinus and BCA (**Figure 2a and 3c**). However, only the SR-BI^ΔCT/ΔCT^/*Ldlr^−/−^* mice formed plaques in coronary arteries (**Figure 2b-d and 3b**). In addition, WD-fed SR-BI^ΔCT/ΔCT^/*Ldlr^−/−^*mice spontaneously developed MI (12 out of 16 mice, **Figure 2f-h,k**) in the right ventricles, left ventricles and apical regions of their hearts as evidenced both macroscopically by cardiac blanching and histologically by the presence of collagen deposition, indicative of fibrotic infarcts, which were not present in the hearts of WD-fed SR-BI^WT/WT^/*Ldlr^−/−^* mice (**Figure 2e**). Moreover, atrial fibrosis was also seen in a subset of WD-fed SR-BI^ΔCT/ΔCT^/*Ldlr^−/−^*mice (**Figure 2i,j**). The distribution and approximate locations of MI and coronary atherosclerosis (**Figure 2k,l**) highlight the spontaneous nature of the model which better represents the diversity of human pathology compared to surgical models of MI.

**Figure 2:**
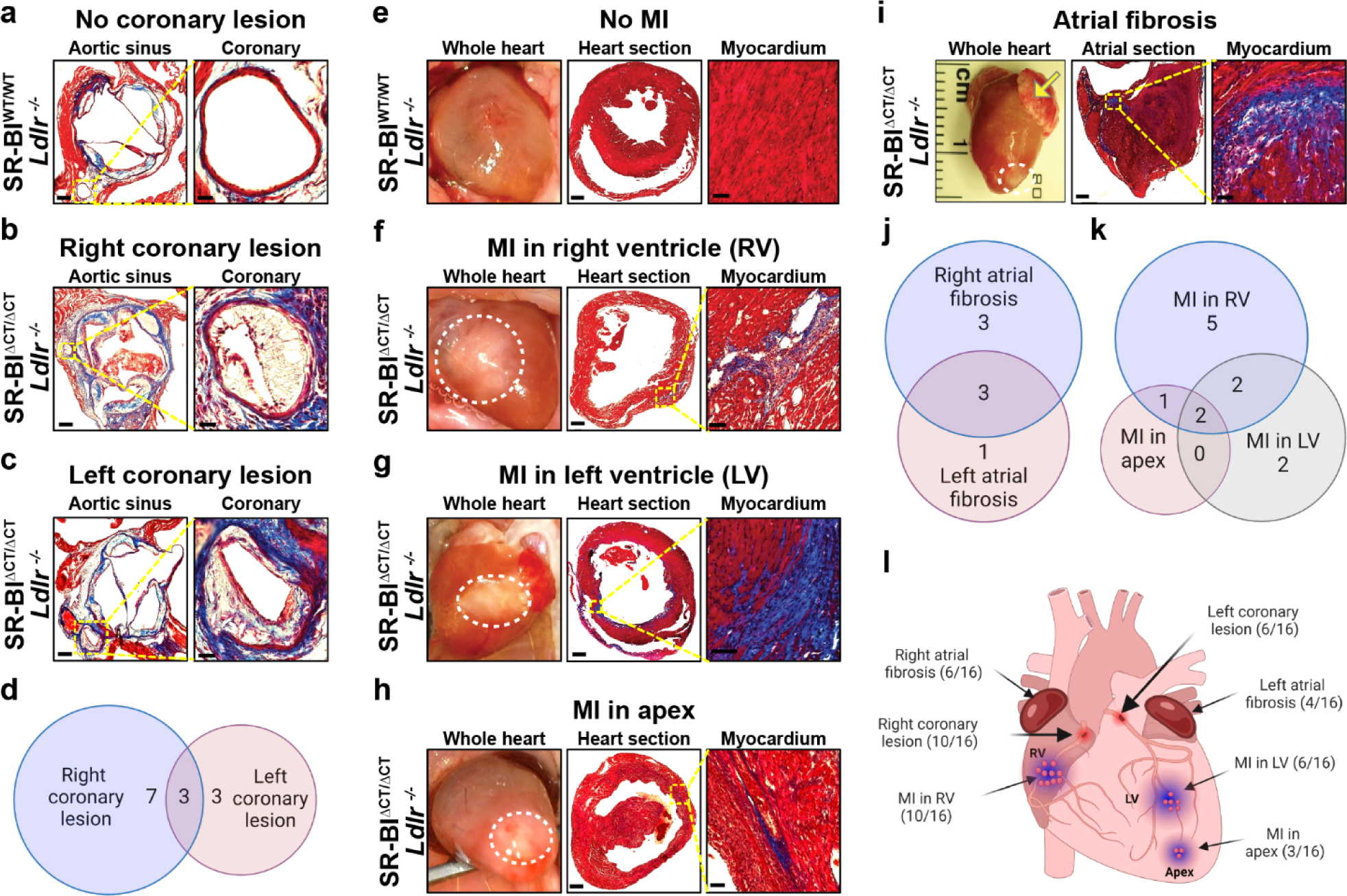
SR-BI^ΔCT/ΔCT^/*Ldlr^−/−^* mice fed a Western Diet develop coronary atherosclerosis and spontaneous myocardial infarction (MI). **(a-c)** Representative images of Masson’s trichrome stained sections of aortic sinus (left, scale bar = 200 µm) and coronary artery (yellow dashed box magnified on the right, scale bar = 50 µm) from SR-BI^WT/WT^/*Ldlr^−/−^* mice (n = 15) (**a**) and SR-BI^ΔCT/ΔCT^/*Ldlr^−/−^* mice (n = 16) (**b,c**) fed a Western Diet (WD) for 26 weeks. Left coronary arteries are shown in **a** and **c** and a right coronary artery is shown in **b**. **(d)** Venn diagram showing co-occurrence of right and left coronary atherosclerotic lesions in WD-fed SR-BI^ΔCT/ΔCT^/*Ldlr^−/−^* mice (n = 16). **(e-h)** Representative mages of whole, excised hearts (left) and Masson’s trichrome stained heart sections at low (middle, scale bar = 200 µm), and high magnification (right, scale bar = 50 µm) from WD-fed mice: **(e)** control SR-BI^WT/WT^/*Ldlr^−/−^* mice (n = 15) showing no evidence of MI, and **(f-h)** SR-BI^ΔCT/ΔCT^/*Ldlr^−/−^* mice (n = 16) showing evidence of MI in right ventricle (RV) **(f)**, left ventricle (LV) **(g),** and apex **(h)**. The white dashed circles highlight blanching that is indicative of myocardial infarcts. Masson’s trichrome stains healthy myocardium red and fibrotic infarcted tissue blue. **(i)** Representative images of a whole, excised heart (left) and Masson’s trichrome stained heart atrial sections at low (middle, scale bar = 200 µm), and high magnification (right, scale bar = 50 µm) from 26 week, WD-fed SR-BI^ΔCT/ΔCT^/*Ldlr^−/−^* mice (n = 16). The yellow arrow indicates fibrosis in the enlarged left atrium and the white dashed circle ndicates myocardial infarct at the apex of the heart. **(j)** Venn diagram showing co-occurrence of right and left atrial fibrosis in SR-BI^ΔCT/ΔCT^/*Ldlr^−/−^* mice (n = 16) fed WD for 26 weeks. **(k)** Venn diagram showing co-occurrence of MI in RV, LV, and apex of SR-BI^ΔCT/ΔCT^/*Ldlr^−/−^* mice (n = 16) fed a WD. **(l)** Cartoon of the heart showing sites of MI observed in SR-BI^ΔCT/ΔCT^/*Ldlr^−/−^* mice (n = 16) fed a WD. Panels **d**, **j**, **k**, and **l** were generated using BioRender.com.

**Figure 3:**
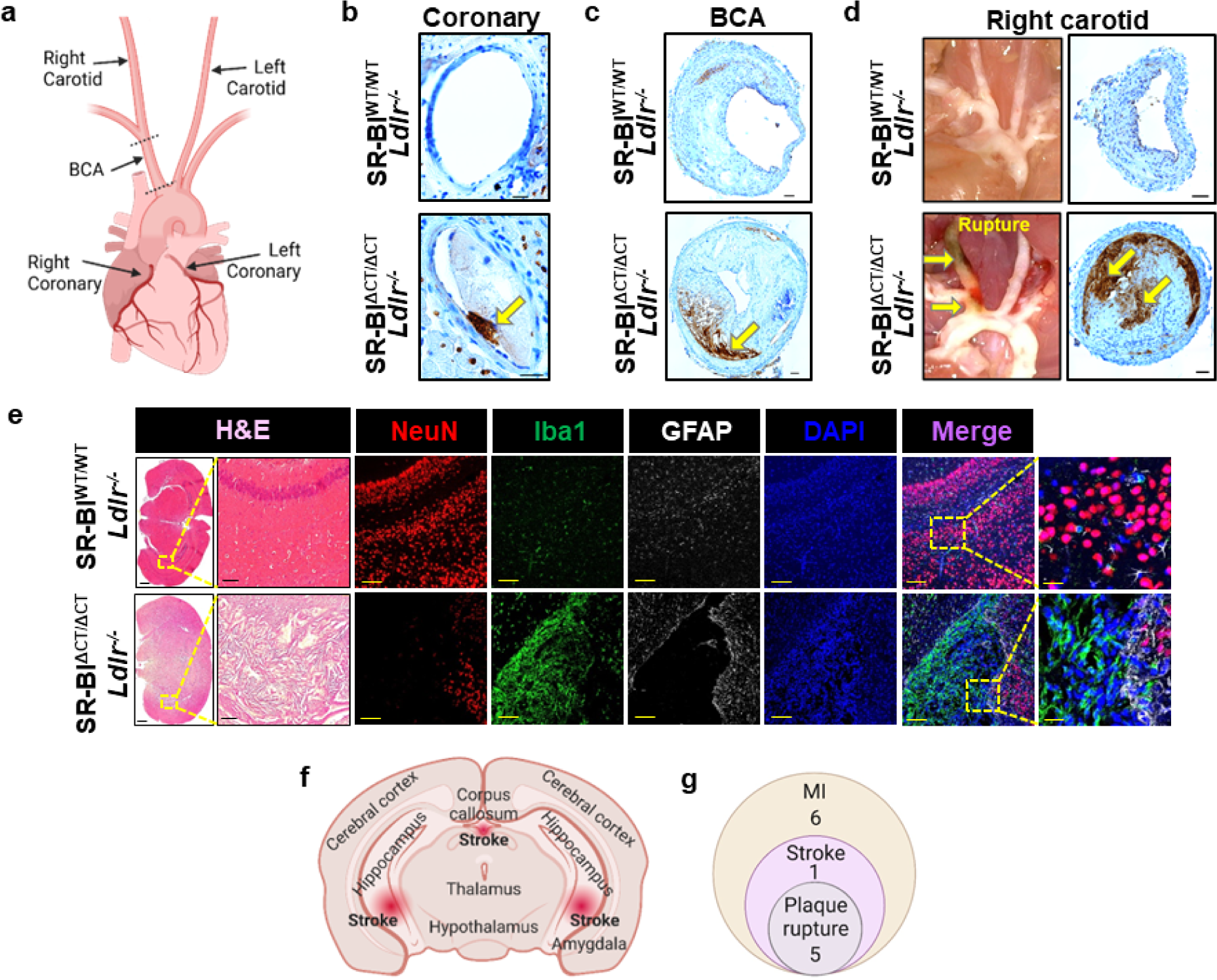
SR-BI^ΔCT/ΔCT^/*Ldlr^−/−^* mice fed a Western Diet exhibit evidence of spontaneous plaque rupture and ischemic stroke. **(a)** Cartoon of the heart and its associated vasculature, including coronary, carotid, and brachiocephalic (BCA) arteries. **(b-d)** Sections of coronary arteries (scale bar = 20 µm) **(b**), BCA (scale bar = 50 µm) (**c**) and right carotid artery (scale bar = 50 µm) (**d**, right) from control SR-BI^WT/WT^/*Ldlr^−/−^*(n = 15) (top) and SR-BI^ΔCT/ΔCT^/*Ldlr^−/−^* (n = 16) (bottom) mice fed the WD for 26 weeks and stained with an erythroid cell specific antibody, Ter-119, that detects plaque rupture/intraplaque hemorrhage (brown staining, yellow arrows). In addition, (**d**) (left) shows *in situ* photographs of the carotid arteries and the BCA from above, with plaque rupture in the SR-BI^ΔCT/ΔCT^/*Ldlr^−/−^* (bottom) but not the SR-BI^WT/WT^/*Ldlr^−/−^* (top) mice. **(e)** Sections of brains from control SR-BI^WT/WT^/*Ldlr^−/−^* (n = 15) (top) and SR-BI^ΔCT/ΔCT^/*Ldlr^−/−^* (n = 16) (bottom) mice fed the WD for 26 weeks. The left two panels show low (scale bars = 500 µm) and high magnification (scale bars = 50 µm) staining with H&E. Adjacent panels show immunofluorescence staining using antibodies to the pan neuronal nuclear marker NeuN, the microglial marker Iba1 and the astrocyte marker GFAP, and fluorescence staining with DAPI (nuclei) (scale bars = 50 µm). The right most panels show high magnification merged fluorescence images (scale bars = 20 µm). **(f)** Cartoon of the brain and locations of areas of ischemic stroke observed in the SR-BI^ΔCT/ΔCT^/*Ldlr^−/−^* mice (n = 16). **(g)** Venn diagram indicating co-occurrence of MI, ischemic stroke, and plaque rupture in SR-BI^ΔCT/ΔCT^/*Ldlr^−/−^* WD-fed mice (n = 16). Panels **a, f** and **g** were generated using BioRender.com.

WD-fed SR-BI^ΔCT/ΔCT^/*Ldlr^−/−^* mice (5 out of 16 mice, **Figure 1e**) displayed gross morphological signs of spontaneous plaque rupture in carotid arteries as well as signs of plaque rupture or intraplaque hemorrhage in the BCA and coronary arteries. This was confirmed by the presence of robust staining with the erythroid cell specific antibody Ter-119, which was not detected in WD-fed SR-BI^WT/WT^/*Ldlr^−/−^* littermate controls (**Figure 3a-d**). Additionally, unlike WD-fed SR-BI^WT/WT^/*Ldlr^−/−^* mice, SR-BI^ΔCT/ΔCT^/*Ldlr^−/−^* mice spontaneously developed ischemic stroke (6 of 16 mice, **Figure 1e**) as evidenced by tissue damage concomitant with a decrease in the pan neuronal marker NeuN (neuronal nuclei)^34^ and a concomitant increase in the microglial marker Iba1 (ionized calcium-binding adaptor molecule-1)^34^ and the astrocyte marker GFAP (glial fibrillary acidic protein)^34^ in affected areas of the brain, typically in the hippocampal region (**Figure 3e,f**). These mice also displayed neurological symptoms of stroke including head tilt, spinning and hind limb paralysis. Of interest, all mice that displayed signs of plaque rupture also showed evidence of stroke and MI. All mice displaying signs of stroke also had MI. A subset of mice had only MI (**Figure 3g**). Taken together these data clearly demonstrate that atherogenic WD-fed SR-BI^ΔCT/ΔCT^/*Ldlr^−/−^* mice develop spontaneous advanced coronary artery atherosclerosis concomitant with MI, plaque rupture, and stroke. It is likely that the cardiac and brain pathologies both contribute to the high mortality of WD-fed SR-BI^ΔCT/ΔCT^/*Ldlr^−/−^* mice (**Figure 1d**).

### Selection of MPO as a therapeutic target

It is well-established that inflammation plays a critical role in atherogenesis^35–37^ and there have been multiple Phase 3 clinical trials attempting to use systemic inhibitors of inflammation to augment the beneficial effects of lipid lowering drugs such as statins^38–43^. However, with the exception of colchicine, which inhibits microtubules and is not a selective anti-inflammatory agent, no clinical trials using systemic anti-inflammatories have resulted in FDA approval likely because anti-inflammatories have had modest or no improvement in MACE, instead resulting in a higher incidence of lethal infections^44,45^. The reasons for this are undoubtedly complex. However, it is possible that these therapies inhibit not only the detrimental pro-inflammatory responses, but also the evolutionarily conserved beneficial inflammatory processes. The latter play a critical role in resistance to pathogenic microorganisms, in tissue repair following injury, and in resolution of inflammation. Consistent with this hypothesis, our lab has provided evidence that inhibition of IL-1β signaling in models of advanced atherosclerosis in mice results in a mixture of beneficial and detrimental changes^46^. Thus, as a proof of principle, we sought to use the SR-BI^ΔCT/ΔCT^/*Ldlr^−/−^* mice to identify a more nuanced therapeutic approach that selectively targets detrimental, chronic inflammatory pathways but leaves beneficial acute inflammatory processes intact.

We considered multiple candidate therapeutic agents and chose a myeloperoxidase (MPO) inhibitor. MPO is an enzyme abundantly expressed by neutrophils and plays an important role in innate microbial defenses^3^. Moreover, MPO is abundantly present in human and mouse atherosclerotic lesions^4,5,47^, and its level in the serum or plasma has been shown to predict risk of MACE in patients with chest pain^48^ and acute coronary syndromes^49^. In addition, although intracellular MPO plays a key role in fighting infections, extracellular MPO has been shown to induce tissue oxidation that may lead to plaque destabilization and myocardial damage^6,50^. Consistent with these human data, we found elevated MPO activity in the plasma of WD-fed SR-BI^ΔCT/ΔCT^/*Ldlr^−/−^*mice compared to that of WD-fed SR-BI^WT/WT^/*Ldlr^−/−^*littermate controls (**Figure 4a,b**). In addition, MPO was abundant in the coronary, carotid and BCA lesions of WD-fed SR-BI^ΔCT/ΔCT^/*Ldlr^−/−^* mice, primarily in cholesterol rich necrotic cores and ruptured areas of the lesions, consistent with human pathology^4,5^ (**Figure 4c-e**). We thus hypothesized that treatment of SR-BI^ΔCT/ΔCT^/*Ldlr^−/−^* mice with a selective inhibitor of extracellular MPO would improve survival by reducing the incidence of plaque rupture, MI, and stroke.

**Figure 4:**
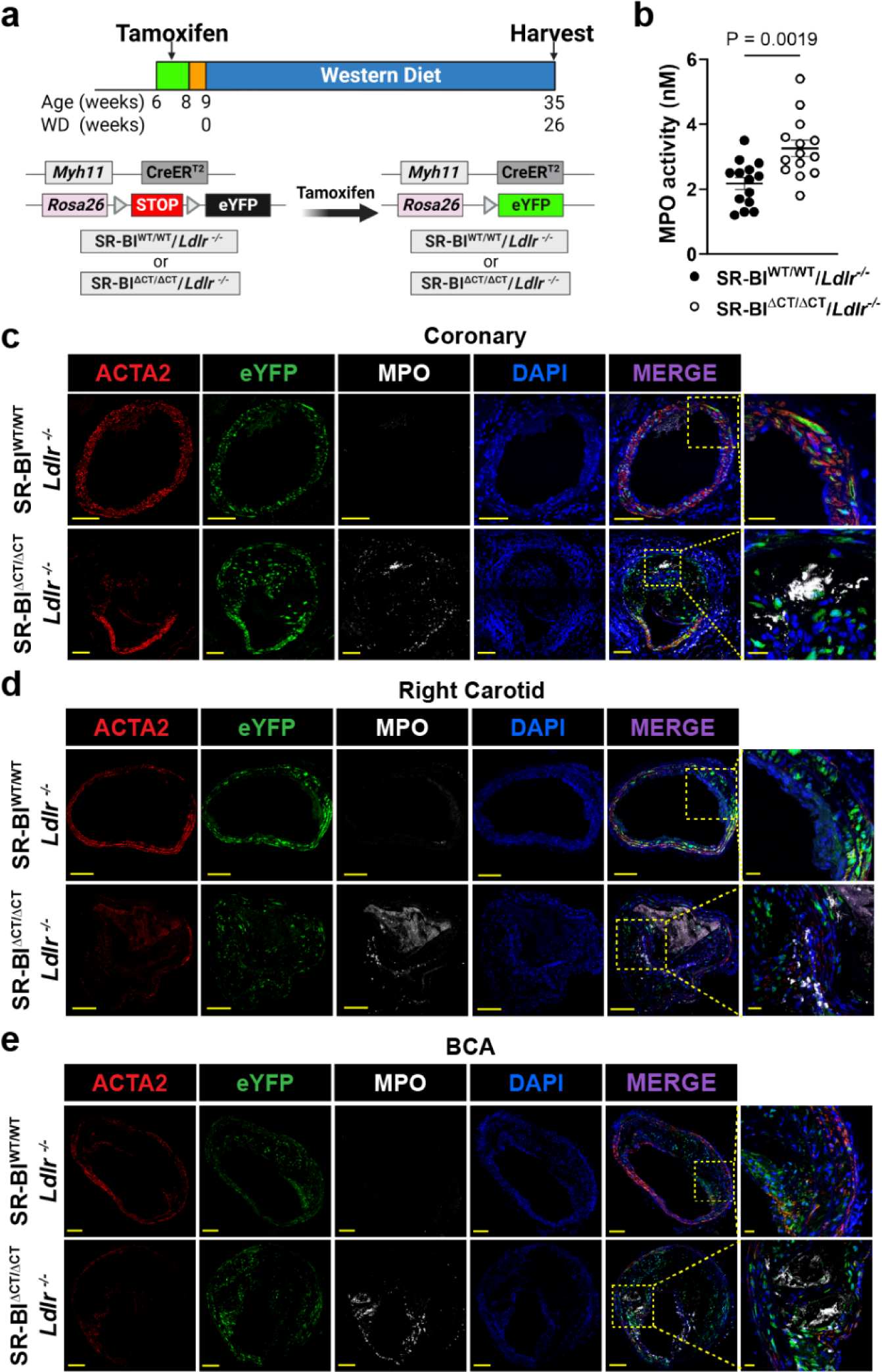
Analysis of MPO levels in the plasma and atherosclerotic lesions of SR-BI^WT/WT^/*Ldlr^−/−^* and SR-BI^ΔCT/ΔCT^/*Ldlr^−/−^* mice fed a Western Diet for 26 weeks. **(a)** Experimental design. **(b)** Plasma MPO enzymatic activity in the plasma of WD-fed SR-BI^WT/WT^/*Ldlr^−/−^* (solid circles, n = 14) and SR-BI^ΔCT/ΔCT^/*Ldlr^−/−^*(open circles, n = 14). Error bars represent mean ± s.e.m. **(c-e)** Representative immunofluorescence images of ACTA2, eYFP (smooth muscle lineage tracing marker), and MPO, and fluorescence of DAPI (nuclei) in coronary (scale bars = 50 µm; merge zoom [far right], scale bars = 20 µm) **(c)**, right carotid artery (scale bars = 100 µm; merge zoom [far right], scale bars = 20 µm) **(d),** and BCA (scale bars = 100 µm; merge zoom [far right], scale bars = 20 µm) **(e)** of SR-BI^WT/WT^/*Ldlr^−/−^*(top, n = 12) and SR-BI^ΔCT/ΔCT^/*Ldlr^−/−^* mice (bottom, n = 14). Statistical analyses were performed using unpaired Student’s *t* test. Panel **a** was generated using BioRender.com.

### Inhibition of MPO reduces MACE

To determine if inhibition of MPO could improve survival and reduce MACE in WD-fed SR-BI^ΔCT/ΔCT^/*Ldlr^−/−^* mice, littermates were fed either a standard WD or WD formulated with the MPO inhibitor AZM198 added (WD+AZM198) for 26 weeks (**Figure 5a**). AZM198 belongs to the class of 2-thioxanthines and is a suicide inhibitor of MPO previously shown to inhibit >90% of extracellular MPO activity without compromising neutrophils’ ability to phagocytose and kill bacteria^7,51,52^. It also has highly acceptable pharmacokinetic and safety profiles^53^. Treatment of WD-fed SR-BI^ΔCT/ΔCT^/*Ldlr^−/−^*mice with AZM198 significantly reduced plasma MPO activity (**Figure 5b**) to that seen in WD-fed SR-BI^WT/WT^/*Ldlr^−/−^* mice (**Figure 4b and dashed line in Figure 5b**). Of major significance, AZM198 treatment of the WD-fed SR-BI^ΔCT/ΔCT^/*Ldlr^−/−^* mice resulted in dramatically improved survival. At 26 weeks after starting the atherogenic diets, 63% (10/16 mice) of the WD+AZM198-fed SR-BI^ΔCT/ΔCT^/*Ldlr^−/−^*mice survived compared to only 6% (1/17 mice) of the SR-BI^ΔCT/ΔCT^/*Ldlr^−/−^* mice fed WD without the drug (**Figure 5c**). In accordance with improved survival, AZM198 treatment reduced the incidence of MI from 77% to 31% (59% decrease), the incidence of stroke from 29% to 13% (57% decrease), and the incidence of plaque rupture from 24% to 6% (73% decrease) based on comparison of WD-fed SR-BI^ΔCT/ΔCT^/*Ldlr^−/−^* mice to WD+AZM198-fed SR-BI^ΔCT/ΔCT^/*Ldlr^−/−^* mice, respectively (**Figure 5d**). Fibrotic area was reduced from 0.27±0.10% in the hearts of WD-fed SR-BI^ΔCT/ΔCT^/*Ldlr^−/−^* mice to 0.03±0.01% in the hearts of WD+AZM198-fed SR-BI^ΔCT/ΔCT^/*Ldlr^−/−^* mice (**Figure 5e**). Moreover, the number of SR-BI^ΔCT/ΔCT^/*Ldlr^−/−^* mice with severely occluded (>50%)^54^ coronary vessels was reduced (90% in WD-fed mice compared to 44% of WD+AZM198-fed mice) as was the average coronary artery lesion size (85.80×10^4^±20.05 µm^2^ for WD-fed mice compared to 26.08×10^4^±4.90 µm^2^ for WD+AZM198-fed mice) (**Figure 5f,g**). The percentage of SR-BI^ΔCT/ΔCT^/*Ldlr^−/−^* mice with plaque rupture/intraplaque hemorrhage, as determined by Ter-119 positive staining, was also reduced in WD+AZM198-fed mice compared to WD-fed littermate controls in coronary (40% in WD vs 8% in WD+AZM198), right carotid (24% in WD vs 6% in WD+AZM198) and BCA lesions (64% in WD vs 17% in WD+AZM198) (**Figure 5h**). Taken together, these data demonstrate that AZM198 inhibited MPO activity concomitant with increased survival and decreased occurrence of MI, stroke, and plaque rupture in WD-fed SR-BI^ΔCT/ΔCT^/*Ldlr^−/−^*mice.

**Figure 5:**
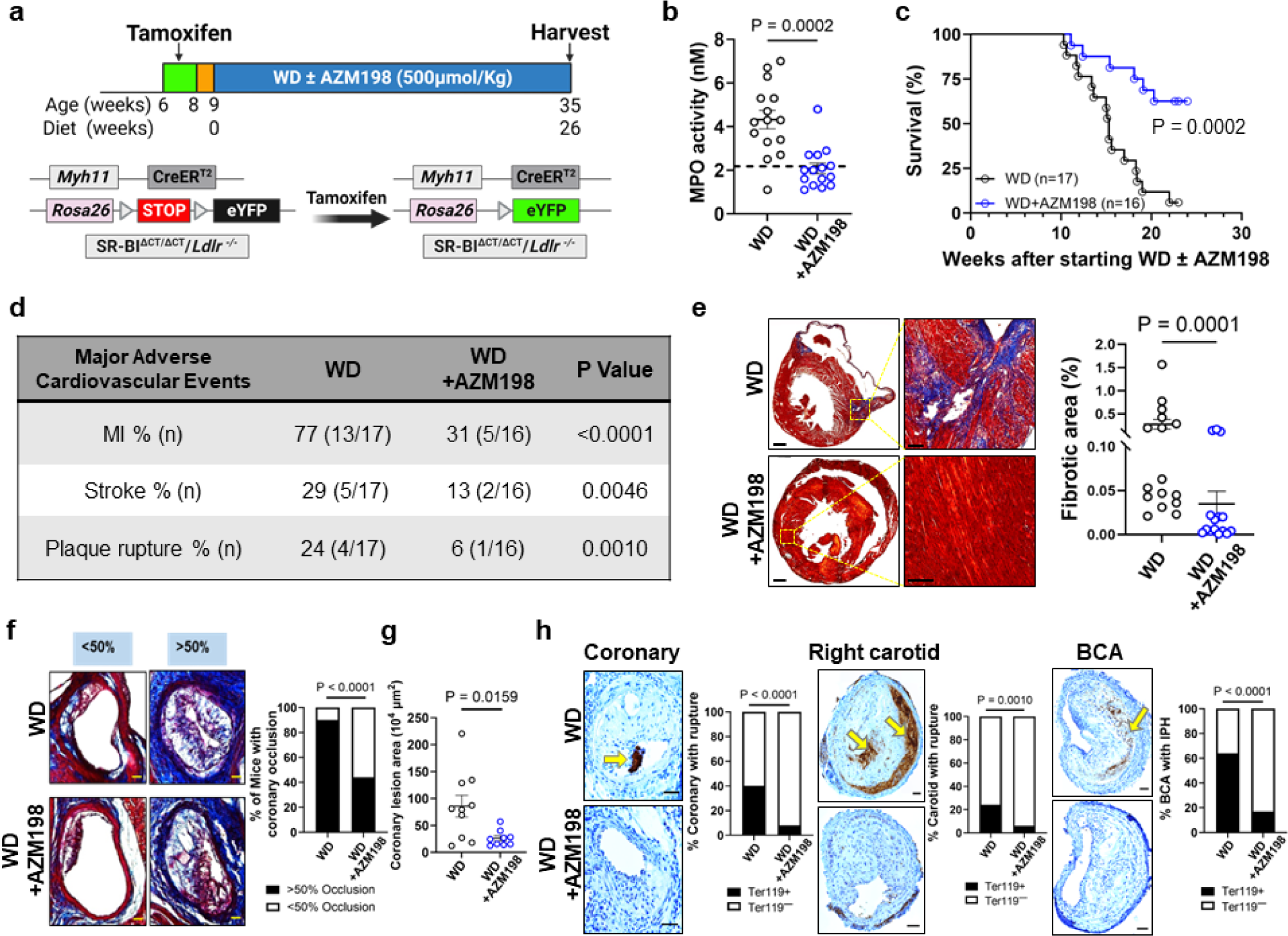
Increased survival and reduced incidence of MI, plaque rupture and ischemic stroke in SR-BI^ΔCT/ΔCT^/*Ldlr^−/−^* mice treated with AZM198. **(a)** Experimental design. **(b)** Plasma MPO enzymatic activity from SR-BI^ΔCT/ΔCT^/*Ldlr^−/−^* mice fed a Western Diet (WD) without (black circles, n = 15) or with (blue circles, n = 15) WD+AZM198 for 26 weeks. **(c)** Kaplan-Meier survival curves (defined by humane terminal endpoint criteria, see Methods) for SR-BI^ΔCT/ΔCT^/*Ldlr^−/−^* mice fed the WD without (black, n = 17) or with (blue, n = 16) AZM198 for 26 weeks. **(d)** Percentage of Major Adverse Cardiovascular Events in SR-BI^ΔCT/ΔCT^/*Ldlr^−/−^* mice fed WD (n = 17) or WD+AZM198 (n = 16) for 26 weeks. **(e)** Representative Masson’s trichrome stained heart sections (left, scale bar = 200 µm; yellow boxed areas magnified in the center, scale bar = 50 µm) and quantitation of fibrotic areas (blue staining) (right panels) from SR-BI^ΔCT/ΔCT^/*Ldlr^−/−^* mice fed the WD without (top or black circles, n = 17) or with (bottom or blue circles, n = 16) AZM198. Healthy myocardium is red and fibrotic tissue is blue. **(f)** Representative Masson’s trichrome stained images of coronary arteries with occlusions filling <50% of the lumen (left) or >50% (center) from SR-BI^ΔCT/ΔCT^/*Ldlr^−/−^* mice fed the WD without (top, n = 10) or with (bottom, n = 9) AZM198. Scale bars = 50 µm. Right graph shows quantitation of the percentage of mice with coronary occlusion filling >50% (black bars) or <50% (white bars) of the lumens. **(g)** Coronary lesion area analysis for SR-BI^ΔCT/ΔCT^/*Ldlr^−/−^* fed a Western Diet (WD) without (black circles, n = 10) or with (blue circles, n = 9) AZM198 for 26 weeks. **(h)** Representative Ter-119 staining of heart sections with coronary arteries and quantification of plaque rupture and intraplaque hemorrhage (IPH) in coronary arteries (left, scale bar = 50 µm), right carotid arteries (center, scale bar = 100 µm) and the BCA (right, scale bar = 100 µm) in SR-BI^ΔCT/ΔCT^/*Ldlr^−/−^* mice fed a Western Diet (WD) without (top and left bars, n = 12) or with (bottom and right bars, n = 14) AZM198 for 26 weeks. The yellow arrows indicate plaque rupture or intraplaque hemorrhage (IPH). Error bars represent mean ± s.e.m. Statistical analyses were performed using unpaired Mann-Whitney *U* test (**b** and **e**), Log-rank (Mantel-Cox) test (**c**), unpaired Student’s *t* test (with Welch’s correction) (**g**) and chi-square (Fisher’s exact) test (**d**, **f**, and **h**). Panel **a** generated using BioRender.com.

### Pleiotropic effects after MPO inhibition

In addition to reduced incidences of MACE (MI, stroke, and plaque rupture), we also found that AZM198 administration had other beneficial effects on WD-fed SR-BI^ΔCT/ΔCT^/*Ldlr^−/−^*mice. There was no difference in food intake between SR-BI^ΔCT/ΔCT^/*Ldlr^−/−^*mice fed WD compared to those fed WD+AZM198 (**Extended Data Figure 4a**). The body weights of the WD-fed mice began to decline after 13 weeks feeding whereas those of mice fed WD+AZM198 were maintained (**Extended Data Figure 4b**). This correlated with WD+AZM198-fed mice having significantly more epididymal fat compared to WD-fed mice (**Extended Data Figure 4c**). As expected, AZM198 was only detected in the plasma from SR-BI^ΔCT/ΔCT^/*Ldlr^−/−^* mice fed the WD+AZM198 diet (**Extended Data Figure 4d**).

Cardiac hypertrophy was reduced in WD+AZM198-fed compared to WD-fed mice whether measured as a ratio normalized to body weight or as absolute weight (1.27±0.10% in WD versus 0.85±0.09% in WD+AZM198 for heart weight/body weight ratio; 0.32±0.02g in WD versus 0.25±0.01g in WD+AZM198.) (**Extended Data Figure 4e**). Longitudinal echocardiography was performed on both the WD- and WD+AZM198-fed SR-BI^ΔCT/ΔCT^/*Ldlr^−/−^* mice to look for overt changes in cardiac function over time (**Extended Data Figure 4f**). Baseline fractional shortening was measured in all mice one day prior to the start of WD feeding. For WD-fed mice (no AZM198), the fractional shortening decreased (indicating reduced heart function) at week 10 of WD feeding and remained low through week 15. Thus, WD-feeding of SR-BI^ΔCT/ΔCT^/*Ldlr^−/−^*mice reduced cardiac function, which likely contributed to their high mortality (**Figure 5c**). Administration of AZM198 to WD-fed SR-BI^ΔCT/ΔCT^/*Ldlr^−/−^*mice did not significantly alter changes in fractional shortening by week 10. However, by week 15 when almost 50% of the WD-fed mice had died, WD+AZM198-fed mice had significantly increased fractional shortening indicating improved heart function which persisted through week 19 (**Extended Data Figure 4f**). Improved cardiac function likely contributed to the increased survival of AZM198 treated mice.

In addition to improving cardiac function, AZM198 markedly decreased liver weight (whether measured as normalized to body weight or absolute weight) and damage in SR-BI^ΔCT/ΔCT^/*Ldlr^−/−^* mice as evidenced by decreased plasma ALT and AST liver enzymes (**Extended Data Figure 5a-c**), reduced liver steatosis as evidenced by reduced Oil Red O staining (**Extended Data Figure 5d,e**), as well as decreased incidence of liver fibrosis as evidenced by reduced Picrosirius Red (PSR)/Fast Green staining (**Extended Data Figure 5f,g**) Of note, it has been previously reported that MPO expression and activity is elevated in both human subjects and experimental mouse models of non-alcoholic steatohepatitis (NASH)^55,56^. Moreover, MPO knockout or pharmacological inhibition of MPO using AZM198 significantly attenuated NASH-induced liver fibrosis and injury^56–58^. It is not clear what role, if any, hepatic pathology plays in the cardiovascular disease in the WD-fed SR-BI^ΔCT/ΔCT^/*Ldlr^−/−^*mice. However, improved hepatic fibrosis in the AZM198 treated mice likely contributed to their improved survival as fibrosis severity strongly predicts mortality in human non-alcoholic fatty liver disease (NAFLD)^59^.

To begin to unravel the mechanisms by which the MPO inhibitor AZM198 increased plaque stability and improved survival rates, we performed analyses of plasma lipids and cytokines, as well as differential leucocyte counts on blood (**Extended Data Figure 6**), and single cell RNA-seq analysis of the BCA lesion areas from WD-fed versus WD+AZM198-fed SR-BI^ΔCT/ΔCT^/*Ldlr^−/−^*mice (**Extended Data Figure 7a**). Analysis of the plasma of WD+AZM198-fed SR-BI^ΔCT/ΔCT^/*Ldlr^−/−^* mice showed no significant differences in total cholesterol or triglycerides between WD-fed and WD+AZM198-fed SR-BI^ΔCT/ΔCT^/*Ldlr^−/−^*mice (**Extended Data Figure 6a**). However, splenomegaly was reduced in the WD+AZM198-fed mice (**Extended Data Figure 6b**). In addition, administration of AZM198 resulted in significant decreases in the concentrations of blood lymphocytes, monocytes, eosinophiles and basophils, as well as reduced plasma concentrations of IL-1β, whereas there were no significant AZM198-induced changes in total white blood cells, neutrophils or other cytokines tested (MIP-1α, MIP-2, TNF-α, IL-6 and IL-10) (**Extended Data Figure 6c,d**). Taken together, the preceding data suggest that the beneficial effects of the MPO inhibitor AZM198 in reducing plaque rupture, MI, and stroke as well as improved liver function, are due at least in part to reduced IL-1β-induced systemic inflammation.

For scRNA-seq analysis, BCA regions (shown in **Extended Data Figure 7a**) from 3 mice per experimental group were pooled together. Thus, these data represent the composite results from multiple replicate mouse samples but with n = 1 per experimental group. As such, our data captures the effective average from replicate mouse samples but lacks the variance information. Consistent with results of previous studies by our lab^31,32^ and others^60^, results from Principal component analysis (PCA), followed by graph-based clustering and Louvain algorithm, delineated 18 transcriptomic clusters comprised of the major cell types found in blood vessels and atherosclerotic plaques (**Extended Figure 7b**). Clusters were classified based on canonical marker gene expression (**Extended Data Figure 8a-i**). Clusters 1-7 are SMC-derived based on detection of the *eyfp* transcript (our SMC lineage tracing marker) (**Extended Data Figure 8c**). Cluster 1 represents contractile SMCs whereas clusters 2-4 represent phenotypically modulated (de-differentiated) SMCs. Clusters 5-7 are SMC-derived extracellular matrix (ECM) producing cells. Of major interest, these data show that WD+AZM198-fed mice have a substantially higher proportion of cells in cluster 7 compared to WD-fed mice (**Extended Figure 7c,d**). Cells in this cluster express ECM genes such as *Fn1 and Col2a1* and may represent SMC-derived cells exhibiting an ECM producing, myofibroblast-like phenotype that are involved in formation and maintenance of the protective fibrous cap^32^ (**Extended Data Figure 7d,e**). Differential gene expression analysis showed an increase in *Col2a1* in cluster 7 in WD+AZM198-fed SR-BI^ΔCT/ΔCT^/*Ldlr^−/−^* mice compared to mice fed WD (**Extended Data Figure 9a**). Paradoxically, cells in cluster 7 also express genes associated with an osteochondrogenic phenotype including *Sox9* and *Omd* and it is possible that these genes may also contribute to the regulation of ECM content and organization in these cells^61–63^. In addition to an increase in cluster 7, we found an increased proportion of cells in clusters 15 and 17 (T-cells) in WD+AZM198-fed SR-BI^ΔCT/ΔCT^/*Ldlr^−/−^* mice compared to mice fed WD (**Extended Figure 7c,d**). Interestingly, cluster 15 contains CD8^+^ T-cells that express both naïve T-cell markers such as *Tcf7* and *Lef1* as well as proliferation markers such *Mki67* which have been previously documented in human atherosclerotic lesions^64^. The cells in cluster 17 appear to be CD4^+^CD8^+^ double positive immature T-cells that express genes such as *Tcf7*, *Lef1*, *Rag1, Ccr9* (**Extended Data Figure 7e**). Similar clusters have been previously reported and they are speculated to be “thymocyte-like” T-cells although their existence and possible function is both controversial and unknown respectively^65^. The potential role of these two T-cell clusters with respect to the benefits produced by AZM198 treatment are unclear and will require additional studies. Cluster 18 appears to represent cytotoxic and/or exhausted T-cells expressing markers such as *Tox*, *Pdcd1* and *Gzmk* and these cells were equally represented in BCA samples from both WD and WD+AZM198-fed mice (**Extended Data Figure 7d,e**). Lastly, there was a decreased proportion of cells in clusters 11, 12, 13 (macrophages) in WD+AZM198-fed SR-BI^ΔCT/ΔCT^/*Ldlr^−/−^* mice compared to mice fed WD (**Extended Figure 7c,d**) in agreement with our data showing reduced circulating monocytes in mice treated with AZM198 (**Extended Data Figure 6c**). Differential gene expression analysis showed a reduction in the expression of *Il1b* in WD+AZM198-fed SR-BI^ΔCT/ΔCT^/*Ldlr^−/−^* mice compared to mice fed WD (**Extended Data Figure 9b**), in agreement with our data showing reduced circulating IL-1β in the plasma of mice treated with AZM198 (**Extended Data Figure 6d**). Finally, we performed Reactome pathway analysis of significantly expressed genes in unsorted or eYFP^+^ sorted cells from the scRNA-seq data sets. In unsorted cells, the top 10 up-regulated pathways in WD+AZM198 versus WD-fed mice centered around increased matrix and collagen organization and formation as well as increased CD3 and lymphoid cell responses (signaling and cell-cell interactions) (**Extended Data Figure 10a, left panel**). Conversely, in unsorted cells, the top 10 down-regulated pathways in WD+AZM198 versus WD-fed mice centered around oxidation and immune responses including activation of NADPH oxidases, chemokine signaling and complement cascade (**Extended Data Figure 10a, right panel**). In eYFP^+^ sorted cells, the top 10 up-regulated pathways in WD+AZM198 versus WD-fed mice again centered around extracellular matrix organization and formation. Moreover, the top 10 down-regulated pathways in eYFP^+^ sorted cells included collagen degradation and assembly as well as decreased immune cell interaction and activation. In summary, the scRNA-seq data suggest that treating WD-fed SR-BI^ΔCT/ΔCT^/*Ldlr^−/−^*mice with AZM198 results in 1) an increase in SMC-derived cells producing ECM which correlates with increased plaque stability and reduced incidents of plaque rupture/intraplaque hemorrhage, 2) potential unknown benefits derived from increased T-cell accumulation, and 3) decreased macrophage content and IL-1β signaling which likely result in decreased inflammation in the plaques of AZM198 treated mice. Taken together, these data provide insight into possible mechanisms by which AZM198 elicits its beneficial effects thereby reducing major adverse cardiovascular events in atherogenic WD-fed SR-BI^ΔCT/ΔCT^/*Ldlr^−/−^* mice.

## Discussion

We have developed and characterized a new mouse model for fatal, occlusive atherosclerotic coronary heart disease (SR-BI^ΔCT/ΔCT^/*Ldlr^−/−^*) that develops plaque rupture, MI and stroke in response to feeding an atherogenic Western Diet. We used these mice to test the therapeutic benefits of the oral MPO inhibitor AZM198. AZM198 administration markedly improved survival of WD-fed mice due at least in part to reduced plaque rupture/intraplaque hemorrhage, MI, and stroke. The drug also had multiple other beneficial effects including reducing liver steatosis and fibrosis, improved cardiac function, decreased splenomegaly, and reduced circulating monocytes and IL-1β that all likely contributed to improved survival. While it seems likely that these effects are mediated through the reduction in MPO activity seen with AZM198 treatment (**Figure 5b**), we cannot rule out that AZM198 may have unknown off-target effects that effected disease progression in the WD-fed SR-BI^ΔCT/ΔCT^/*Ldlr^−/−^* mice.

Our findings are consistent with previous studies by Rashid *et al.*^7^ who used the tandem stenosis model and showed that global knockout of MPO or inhibition of MPO with AZM198 between 6 - 19 weeks of age of WD-fed *Apoe^−/−^* mice resulted in a near doubling of the fibrous cap thickness of tandem stenosed artery lesions, which is consistent with increased plaque stability. Similarly, using the same tandem stenosis model in *Apoe^−/−^* mice, an intervention study by Chen *et al.* recently demonstrated that treatment with AZM198 for either 1 week or 5 weeks increased fibrous cap thickness of existing tandem stenosed artery lesions^51^. These important studies employed a surgical technique to induce stenosis (without MI or stroke) and did not establish if AZM198 treatment reduces mortality^51^.

The Likelihood of Approval (LOA) rate for cardiovascular disease drugs is among the lowest of any major disease^66^, whereas the cost of clinical trials is extraordinarily high due to the need to study thousands of patients over multiple years. For example, the CANTOS phase 3 clinical trial of the anti-IL-1β antibody canakinumab included over 10,000 subjects studied over six years at an estimated cost of greater than $776M^38^. The WD-fed SR-BI^ΔCT/ΔCT^/*Ldlr^−/−^* mice are an attractive pre-clinical model of late-stage, diet-inducible atherosclerotic disease that will likely contribute to: 1) our understanding of mechanisms that control plaque stability, 2) identification of novel therapeutic approaches for treating patients with advanced atherosclerosis, and 3) improved LOA rate of human clinical trials aimed at identifying therapies that augment the benefits of lipid lowering drugs. We speculate that the success of the extracellular MPO inhibitor AZM198 used in our studies provides proof of principle of the utility of selective anti-inflammatory agents that target detrimental inflammation without compromising innate immunity, although definitively testing this will require additional studies. This is particularly important given our pandemic susceptible world and the need to administer atherosclerosis-protective drugs for long periods of time.

In summary the SR-BI^ΔCT/ΔCT^/*Ldlr^−/−^* mouse is a novel model of coronary atherosclerosis that spontaneously develop plaque rupture, MI, and stroke, as well as increased mortality in response to Western Diet feeding. We show proof of principle that this mouse can be used to test potential new therapeutics (such as the MPO inhibitor AZM198) to reduce MACE in these mice. This mouse model provides an important tool and will advance the search for druggable targets to treat atherosclerosis.

## Extended Data Figures

**Extended Data Figure 1:**
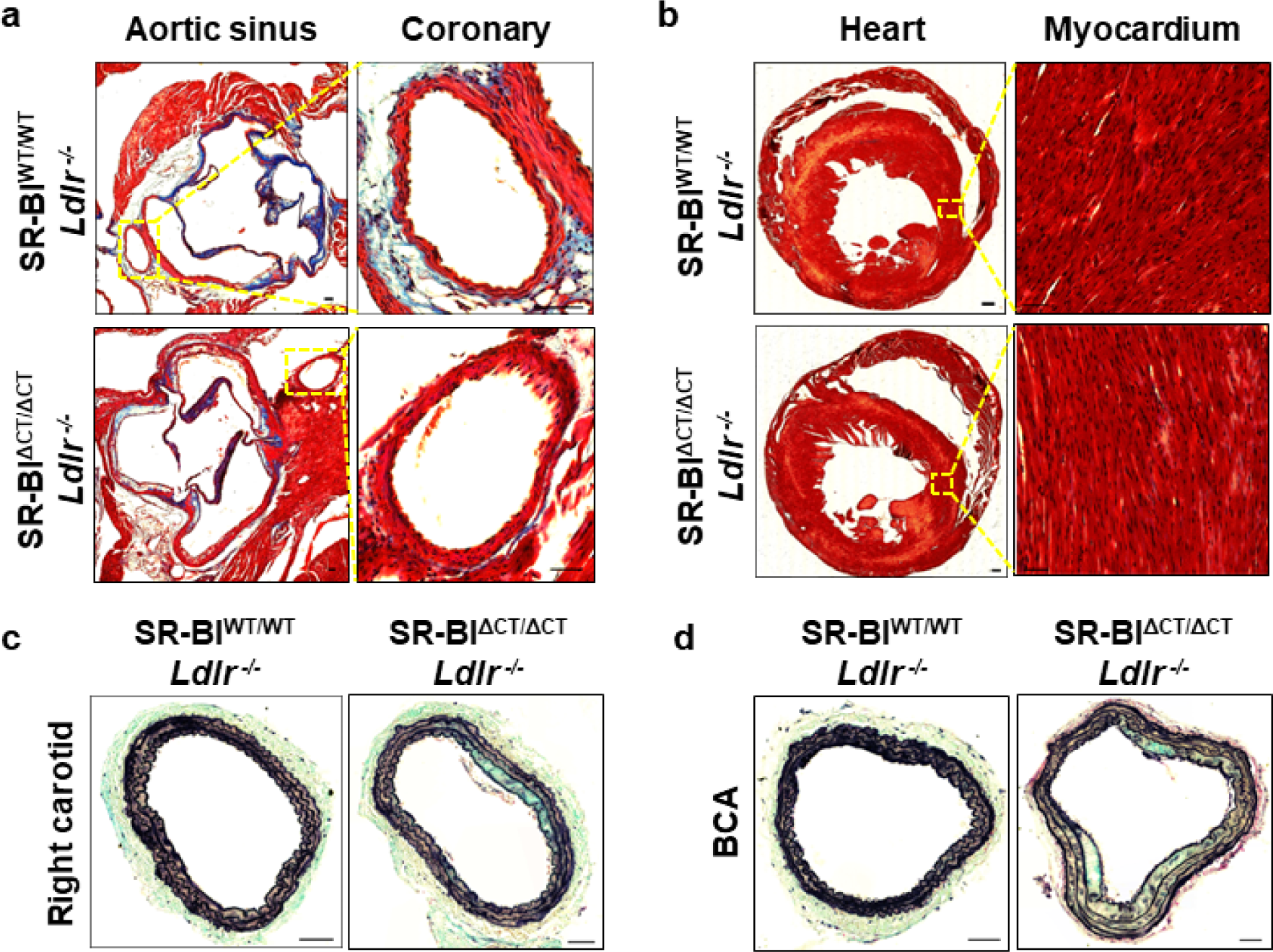
SR-BI^ΔCT/ΔCT^/*Ldlr^−/−^* mice fed a standard rodent Chow diet show no coronary atherosclerosis or spontaneous myocardial infarction (MI). **(a,b)** Representative images of Masson’s trichrome stained sections of aortic sinuses and hearts from SR-BI^WT/WT^/*Ldlr^−/−^*(top, n = 9) and SR-BI^ΔCT/ΔCT^/*Ldlr^−/−^* (bottom, n = 10) mice fed Chow diet for 26 weeks. (**a**) Aortic sinuses (left, scale bar = 200 µm) and coronary artery (yellow box, magnified view on the right, scale bar = 50 µm). (**b**) Heart cross sections (left, scale bar = 500 µm) and myocardium (yellow box, magnified view on he right, scale bar = 50 µm). **(c,d)** Representative Movat pentachrome stained sections of the right carotid artery (**c**) and BCA (**d**) from SR-BI^WT/WT^/*Ldlr^−/−^* (left, n = 9) and SR-BI^ΔCT/ΔCT^/*Ldlr^−/−^* (right, n = 10) mice fed Chow diet for 26 weeks (scale bars = 100 µm).

**Extended Data Figure 2:**
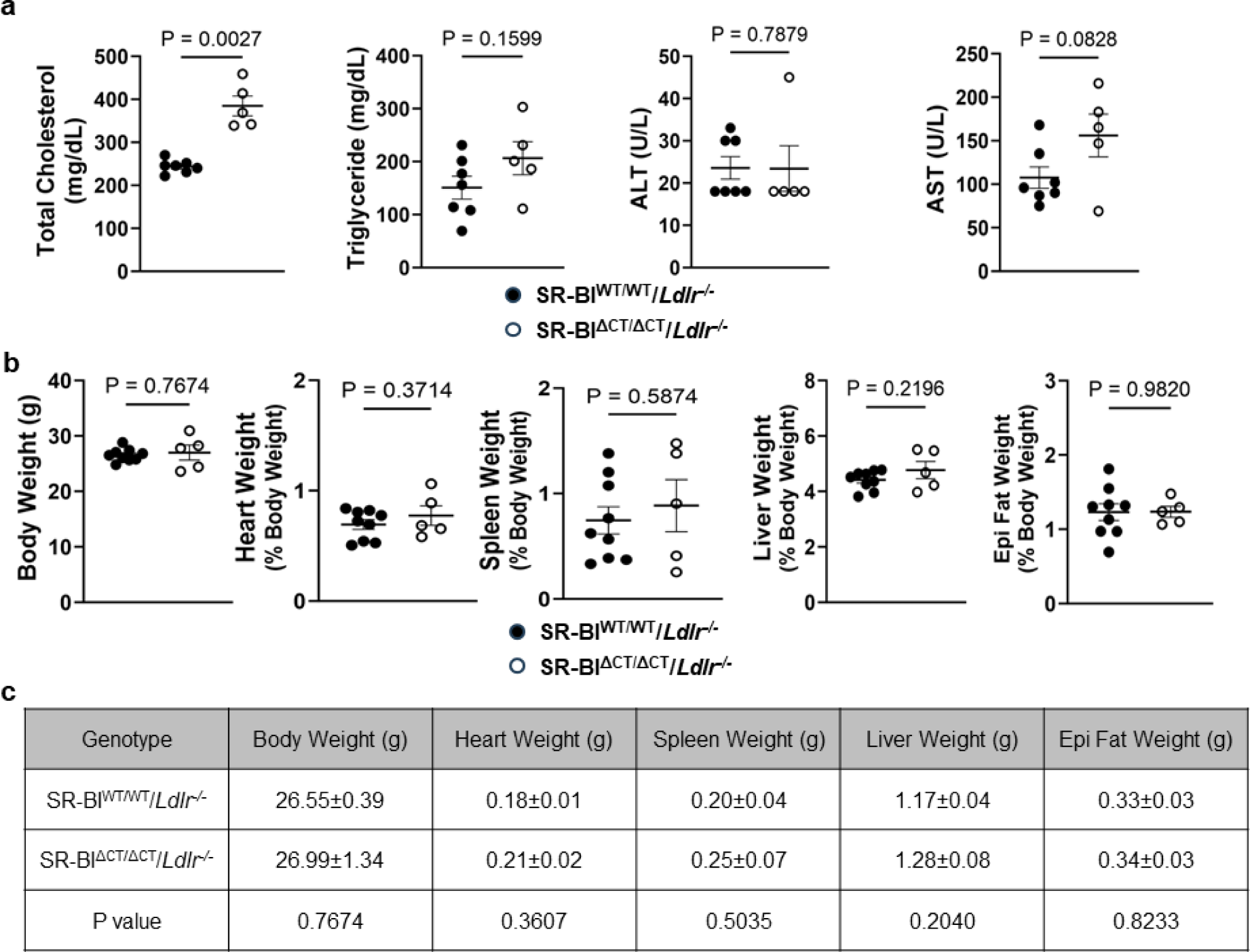
Analysis of plasma and organ weights of SR-BI^WT/WT^/*Ldlr^−/−^* and SR-BI^ΔCT/ΔCT^/*Ldlr^−/−^* mice fed a Chow diet for 26 weeks. **(a)** Plasma levels of total cholesterol, triglycerides, and the liver enzymes alanine transaminase (ALT) and aspartate transaminase (AST) in SR-BI^WT/WT^/*Ldlr^−/−^* (solid circles, n = 9) and SR-BI^ΔCT/ΔCT^/*Ldlr^−/−^* (open circles, n = 5) mice fed a Chow diet for 26 weeks. **(b,c)** Body and heart, spleen, liver and epididymal (epi) fat weights (% of body weight in **b** and absolute weights in **c**) for SR-BI^WT/WT^/*Ldlr^−/−^*(solid circles, n = 9) and SR-BI^ΔCT/ΔCT^/*Ldlr^−/−^* (open circles, n = 5) mice fed a Chow diet for 26 weeks. Data are shown as mean ± s.e.m. Statistical analyses were performed using unpaired Student’s *t* test (with Welch’s correction when variance was unequal) and unpaired Mann-Whitney *U* test.

**Extended Data Figure 3:**
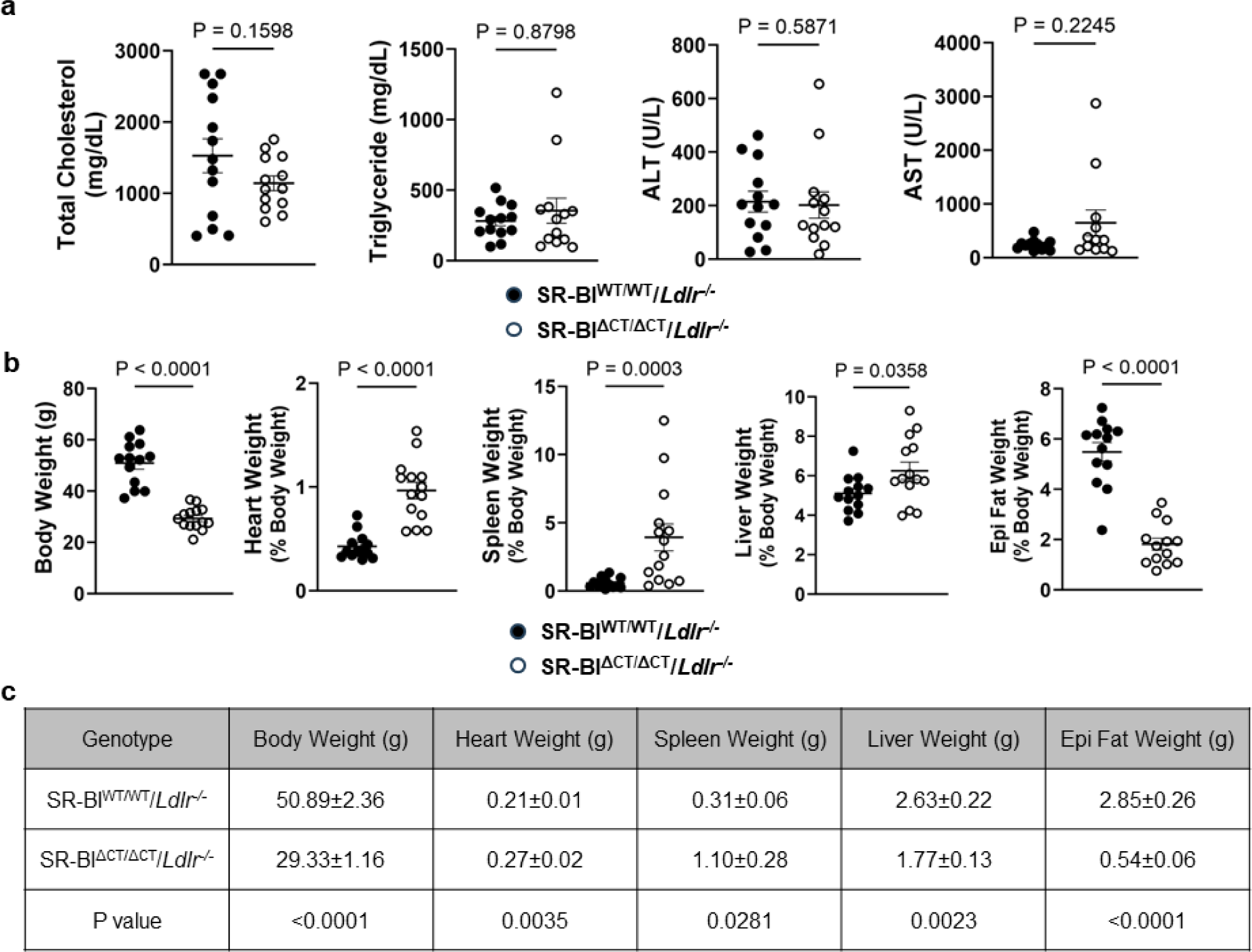
Analysis of plasma and organ weights of SR-BI^WT/WT^/*Ldlr^−/−^* and SR-BI^ΔCT/ΔCT^/*Ldlr^−/−^* mice fed a Western Diet (WD) for 26 weeks. **(a)** Plasma levels of total cholesterol, riglycerides, and the liver enzymes alanine transaminase (ALT) and aspartate transaminase (AST) in SR-BI^WT/WT^/*Ldlr^−/−^* (solid circles, n = 13) and SR-BI^ΔCT/ΔCT^/*Ldlr^−/−^*(open circles, n = 14) mice fed a Western diet for 26 weeks. **(b,c)** Body and heart, spleen, liver and epididymal (epi) fat weights (% of body weight in **b** and absolute weights in **c**) for SR-BI^WT/WT^/*Ldlr^−/−^* (solid circles, n = 13) and SR-BI^ΔCT/ΔCT^/*Ldlr^−/−^*(open circles, n = 14) mice fed a Western diet for 26 weeks. Data are shown as mean ± s.e.m. Statistical analyses were performed using unpaired Student’s *t* test (with Welch’s correction when variance was unequal) and unpaired Mann-Whitney *U* test.

**Extended Data Figure 4:**
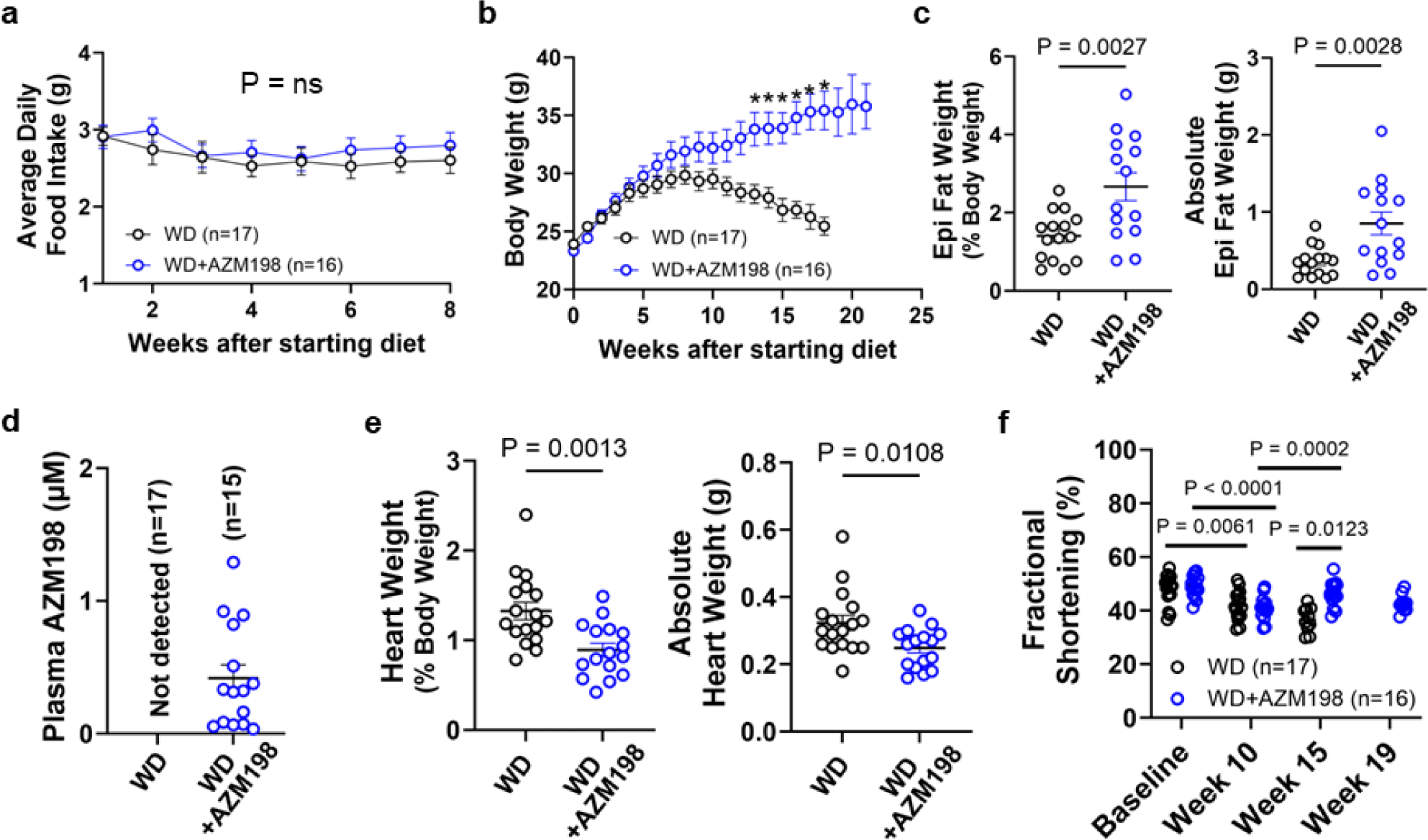
AZM198 improved heart function in SR-BI^ΔCT/ΔCT^/*Ldlr^−/−^* mice fed a Western Diet. SR-BI^ΔCT/ΔCT^/*Ldlr^−/−^* mice were fed a Western Diet (WD) without (black circles) or with AZM198 (blue circles) for the indicated times. (**a**) Average daily food intake (WD, n = 17; WD+AZM198, n = 16). **(b)** Body weights (WD, n = 17; WD+AZM198, n = 16). *P < 0.05. (**c**) Epididymal (Epi) fat weights (% of body weight [left] and absolute weights [right]) (WD, n = 15; WD+AZM198, n = 14). **(d)** Plasma AZM198 levels (WD, n = 17; WD+AZM198, n = 15) **(e)** Heart weights (% of body weight [left] and absolute weights [right]) (WD, n = 17; WD+AZM198, n = 16). **(f)** Fractional shortening in hearts based on sequential echocardiographic analysis (WD, n = 17; WD+AZM198, n = 16). Error bars represent mean ± s.e.m. Statistical analyses were performed using two-way ANOVA with Sidak’s multiple comparisons tests (**a**), Mann-Whitney *U* with Holm-Sidak’s multiple comparisons tests (**b**), unpaired Student’s *t* test (**d, e**) and mixed-effects analysis with Sidak’s multiple comparisons tests (**f**).

**Extended Data Figure 5:**
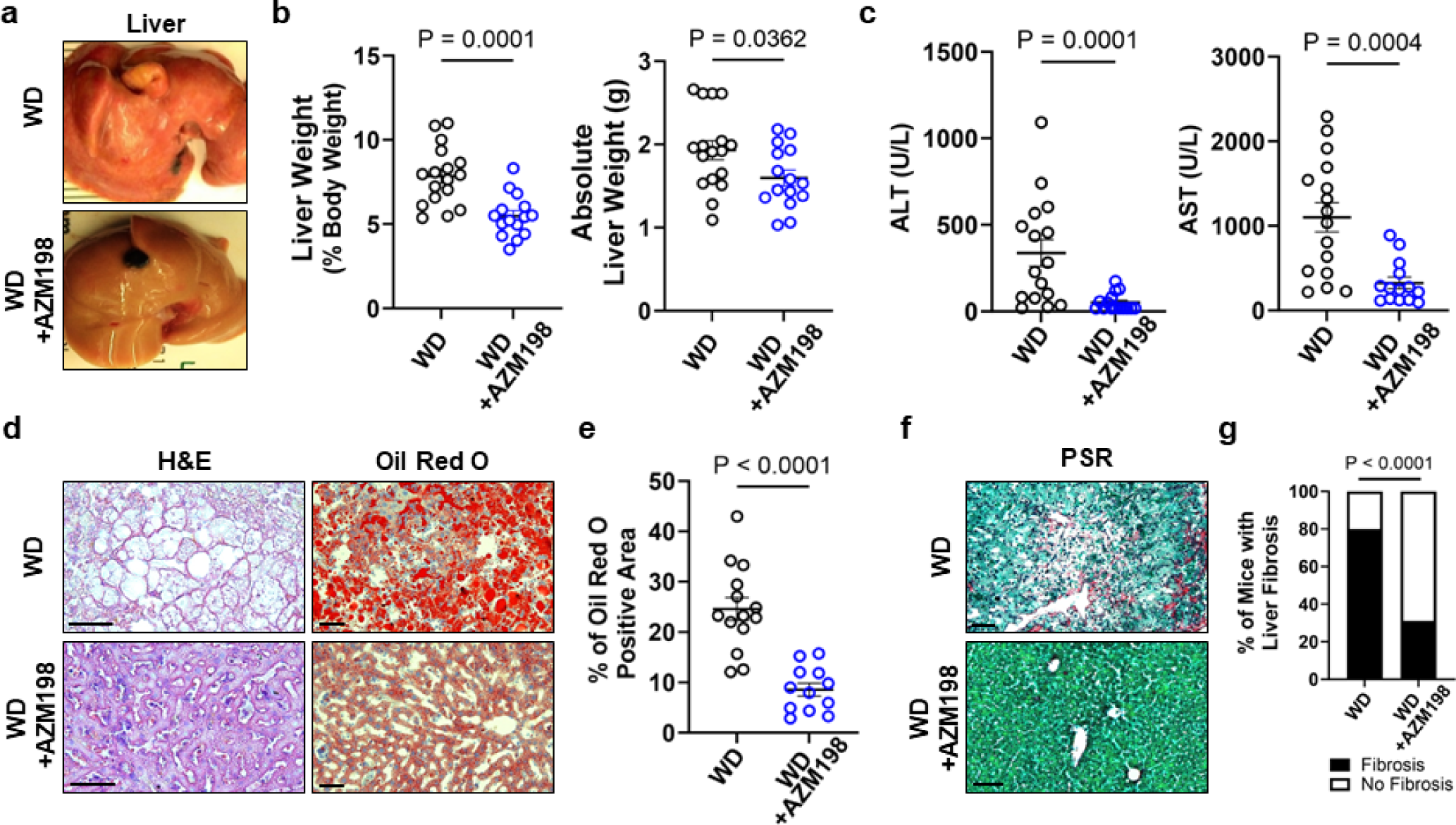
AZM198 markedly improved liver function in SR-BI^ΔCT/ΔCT^/*Ldlr^−/−^* mice fed a Western Diet for 26 weeks. SR-BI^ΔCT/ΔCT^/*Ldlr^−/−^* mice were fed a Western Diet (WD) without (upper panels or black circles) or with AZM198 (lower panels or blue circles) for 26 weeks. **(a)** Representative images of excised livers (WD, n = 17; WD+AZM198, n = 16). **(b)** Liver weights (% of body weight [left] and absolute weights [right]) (WD, n = 17; WD+AZM198, n = 15). **(c)** Plasma evels of the liver enzymes alanine transaminase (ALT) and aspartate transaminase (AST) (WD, n = 16; WD+AZM198, n = 15). **(d)** Representative H&E (left) and Oil Red O (right) images of the livers (WD, n = 16; WD+AZM198, n = 15, scale bars = 50 µm). **(e)** Quantification of fat deposition (% Oil Red O staining) in the livers (WD, n = 14; WD+AZM198, n = 12). **(f,g)** Representative Picrosirius Red/Fast green (PSR) staining of liver sections (**f**, scale bars = 50 µm) and incidence of liver fibrosis (PSR positive staining) (WD, n = 14; WD+AZM198, n = 12). Error bars represent mean ± s.e.m. Statistical analyses were performed using unpaired Student’s *t* test (**b**) unpaired Mann-Whitney *U* test (**c** and **e**), chi-square (Fisher’s exact) test (**g**).

**Extended Data Figure 6:**
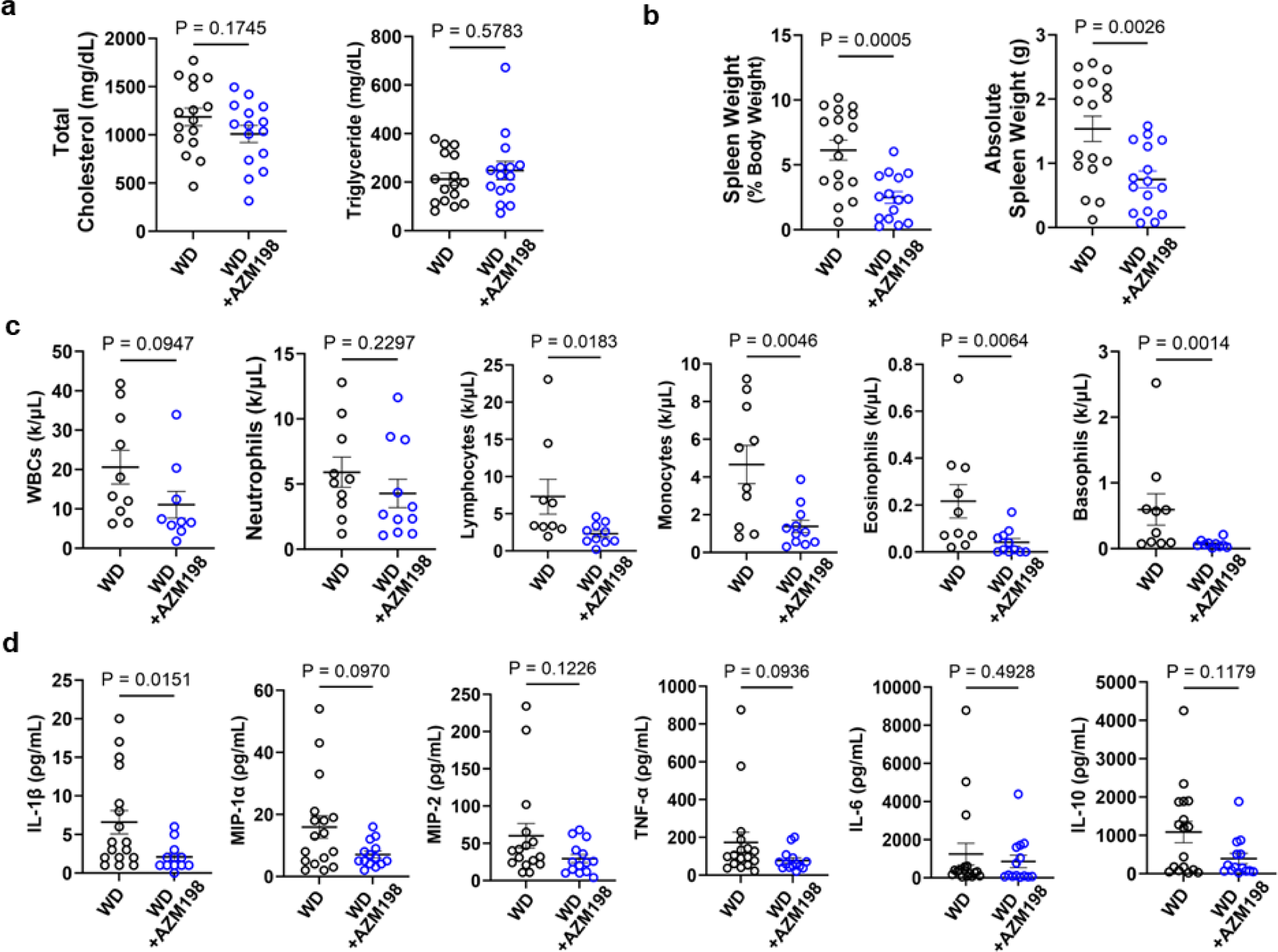
Effects of AZM198 on blood and plasma compositions and spleen weights in SR-BI^ΔCT/ΔCT^/*Ldlr^−/−^* mice fed a Western Diet. SR-BI^ΔCT/ΔCT^/*Ldlr^−/−^*mice were fed a Western Diet (WD) without (black circles) or with AZM198 (blue circles) for 26 weeks. **(a)** Plasma total cholesterol (left) and triglycerides (right) (WD, n = 16; WD+AZM198, n = 16). **(b)** Spleen weights % of body weight [left] and absolute weights [right] (WD, n = 17; WD+AZM198, n = 16). **(c)** Concentrations of blood cells: white blood cells (WBCs), neutrophils, lymphocytes, monocytes, eosinophils and basophils. (WD, n = 10; WD+AZM198, n = 9). **(d)** Cytokine concentrations (WD, n = 16; WD+AZM198, n = 14). Error bars represent mean ± s.e.m. Statistical analyses were performed using unpaired Mann-Whitney *U* test or Student’s *t* test (with Welch’s correction when variance was unequal).

**Extended Data Figure 7:**
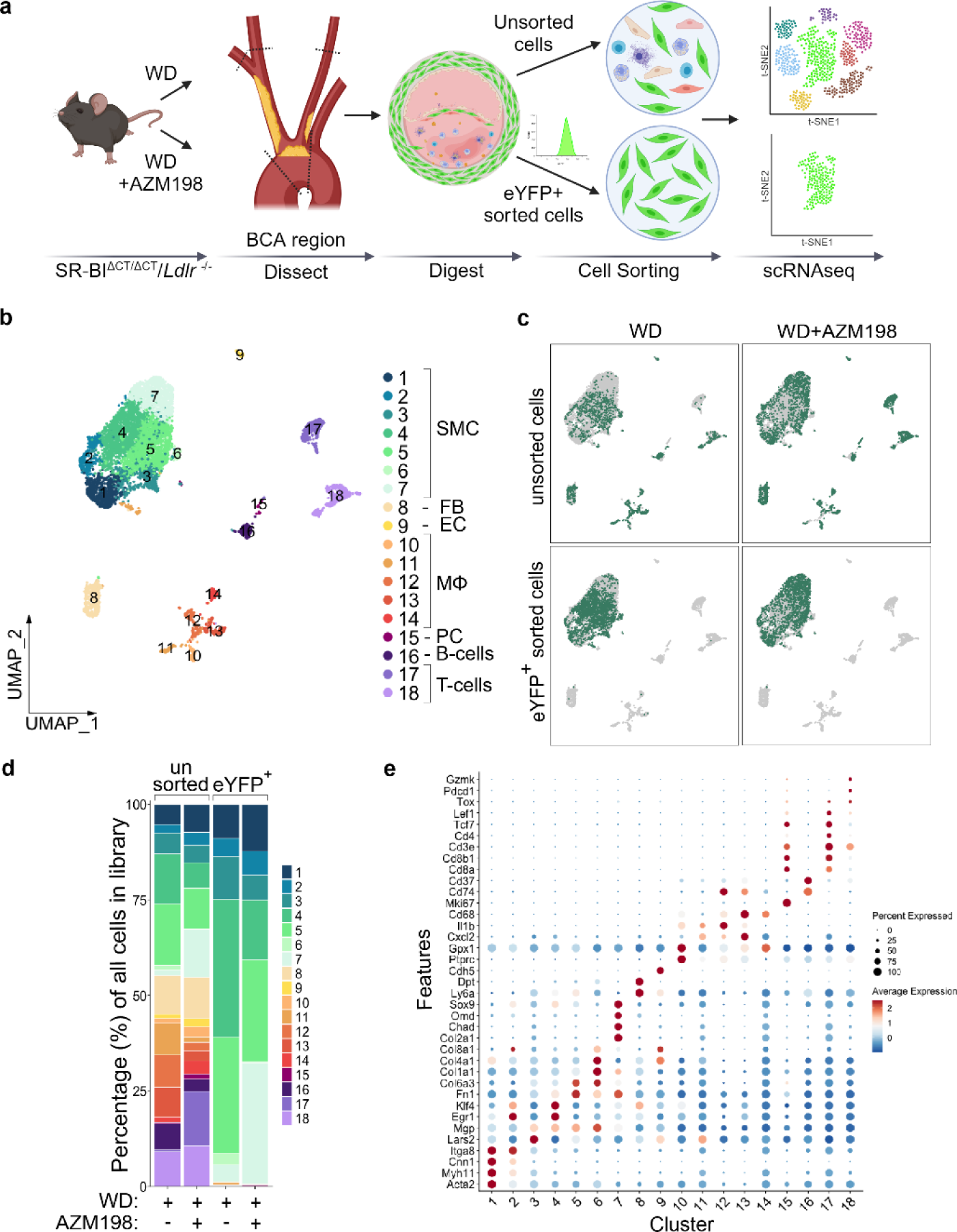
Inhibition of MPO by AZM198 resulted in increased SMC with a myofibroblast-like ECM producing phenotype, decreased macrophages, and increased T cells. **(a)** Experimental design. Samples of BCA regions from three mice for each condition (WD with or without AZM198, fed for 24 weeks) were pooled into one group each and subsequently analyzed. After generation of single cells from the tissue, samples of either the entire population of single cells (unsorted) or eYFP positive cells (SMC cells, eYFP+ sorted cells) were subjected to scRNA-seq analysis. **(b)** UMAP representation of all cells in the scRNA-seq data sets from BCA regions showing the 18 different clusters identified by canonical marker gene expression. **c)** UMAP representation of each individual scRNA-seq library (WD, left; WD+AZM198, right) showing differences in the distribution of cells in each cluster for unsorted (top) and eYFP^+^ sorted (bottom) cells. **(d)** Quantification (%) of unsorted and eYFP^+^ sorted cells in each cluster identified by UMAP analysis for each scRNA-seq data set (WD or WD+AZM198, unsorted and sorted). **(e)** Dotplot analysis showing traditional markers used for cell identification, as well as unique markers related to each cluster. scRNA-seq, single-cell RNA sequencing; SMC, smooth muscle cell; and UMAP, uniform manifold approximation and projection. MPO, myeloperoxidase; eYFP, enhanced yellow fluorescent protein; ECM, extracellular matrix. Panel **a** was generated using BioRender.com.

**Extended Data Figure 8:**
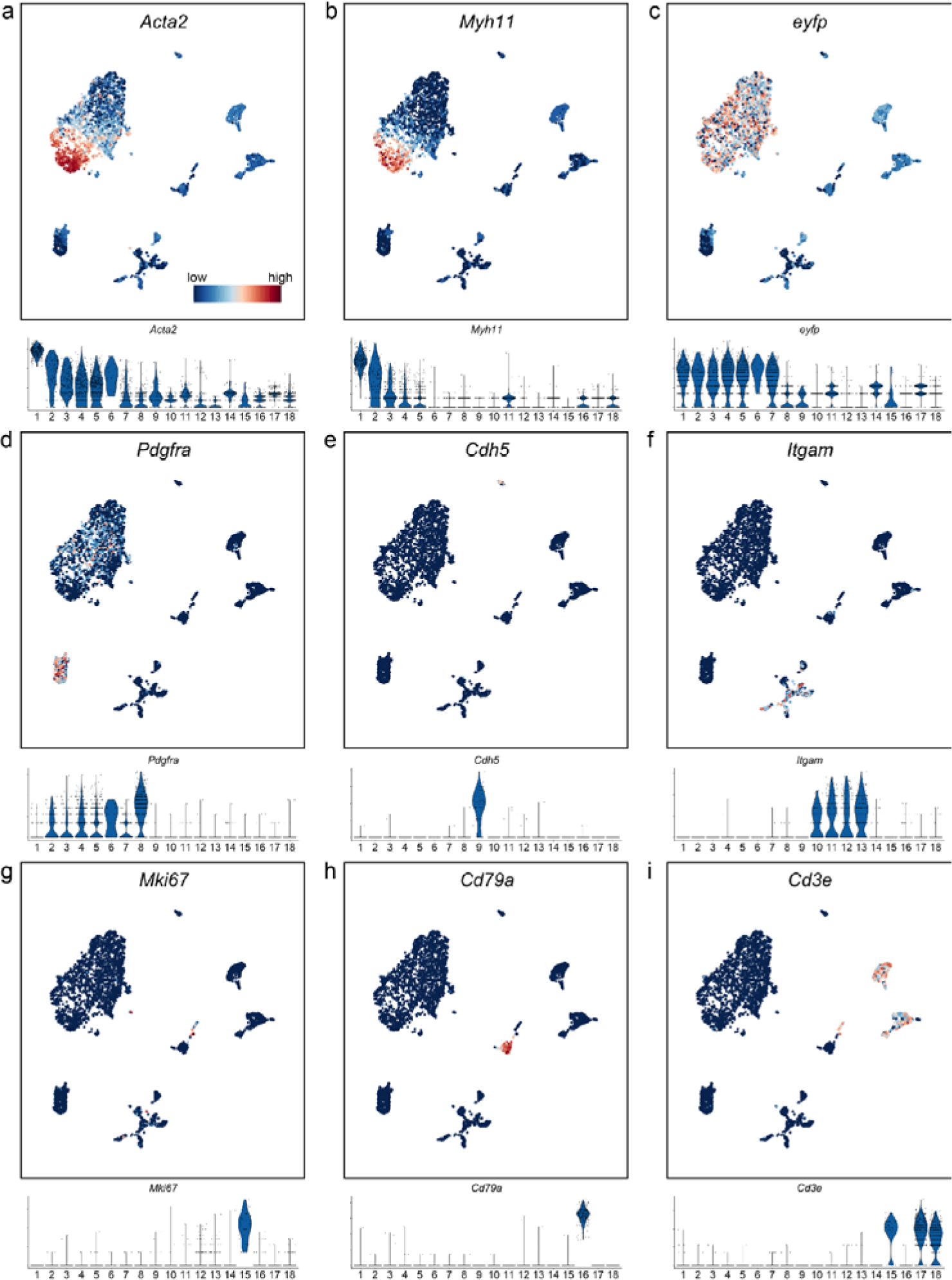
Representative canonical marker gene expression used to identify cell types in each cluster identified by scRNA-seq of BCA regions. Samples of BCA regions from three mice for each condition (WD with or without AZM198) were pooled into one group each and subsequently analyzed. After generation of single cells from the tissue, samples of either the entire population of single cells (unsorted) or eYFP positive cells (SMC cells, eYFP+ sorted cells) were subjected to scRNA-seq analysis. *Acta2* **(a)** and *Myh11* **(b)** are SMC marker genes. *Eyfp* **(c)** denotes all cells derived from SMC lineage. *Pdgfra* **(d)** is a fibroblast marker gene. *Cdh5* **(e)** is an endothelial cell marker gene. *Itgam* **(f)** (also known as *CD11b*) is a macrophage marker gene. *Mki67* **(g)** is a marker gene for proliferating cells. *CD79a* **(h)** is a B-cell marker gene. *Cd3e* **(i)** is a T-cell marker gene.

**Extended Data Figure 9:**
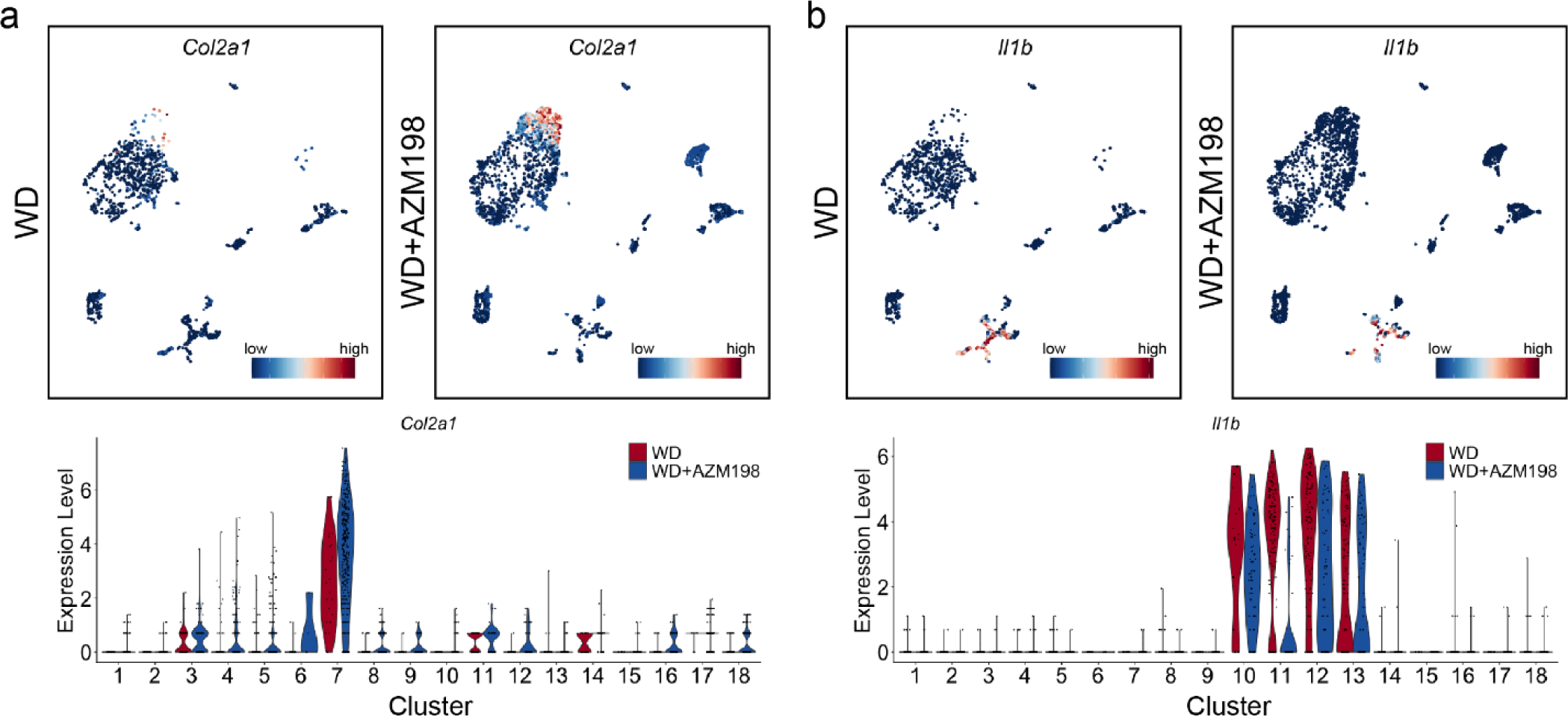
*Col2a1* expression is increased while *Il1b* expression is decreased in SR-BI^ΔCT/ΔCT^/*Ldlr^−/−^* WD+AZM198 versus WD fed mice. Samples of BCA regions from three mice for each condition (WD with or without AZM198) were pooled into one group each and subsequently analyzed. After generation of single cells from the tissue, samples of either the entire population of single cells (unsorted) and eYFP positive cells (SMC cells, eYFP+ sorted cells) were subjected to scRNA-seq analysis. **(a)** UMAP plots showing expression of *Col2a1* all cells isolated from WD (left) and WD+AZM198 fed mice (right). A violin plot shows that the expression of *Col2a1* in cluster 7 is increased in WD+AZM198 fed mice compared to WD fed mice (bottom). **(b)** UMAP plots showing expression of *Il1b* from all cells isolated from WD (left) and WD+AZM198 fed mice (right). A violin plot shows that the expression of *Il1b* in cluster 11 is decreased in WD+AZM198 fed mice compared o WD fed mice (bottom).

**Extended Data Figure 10:**
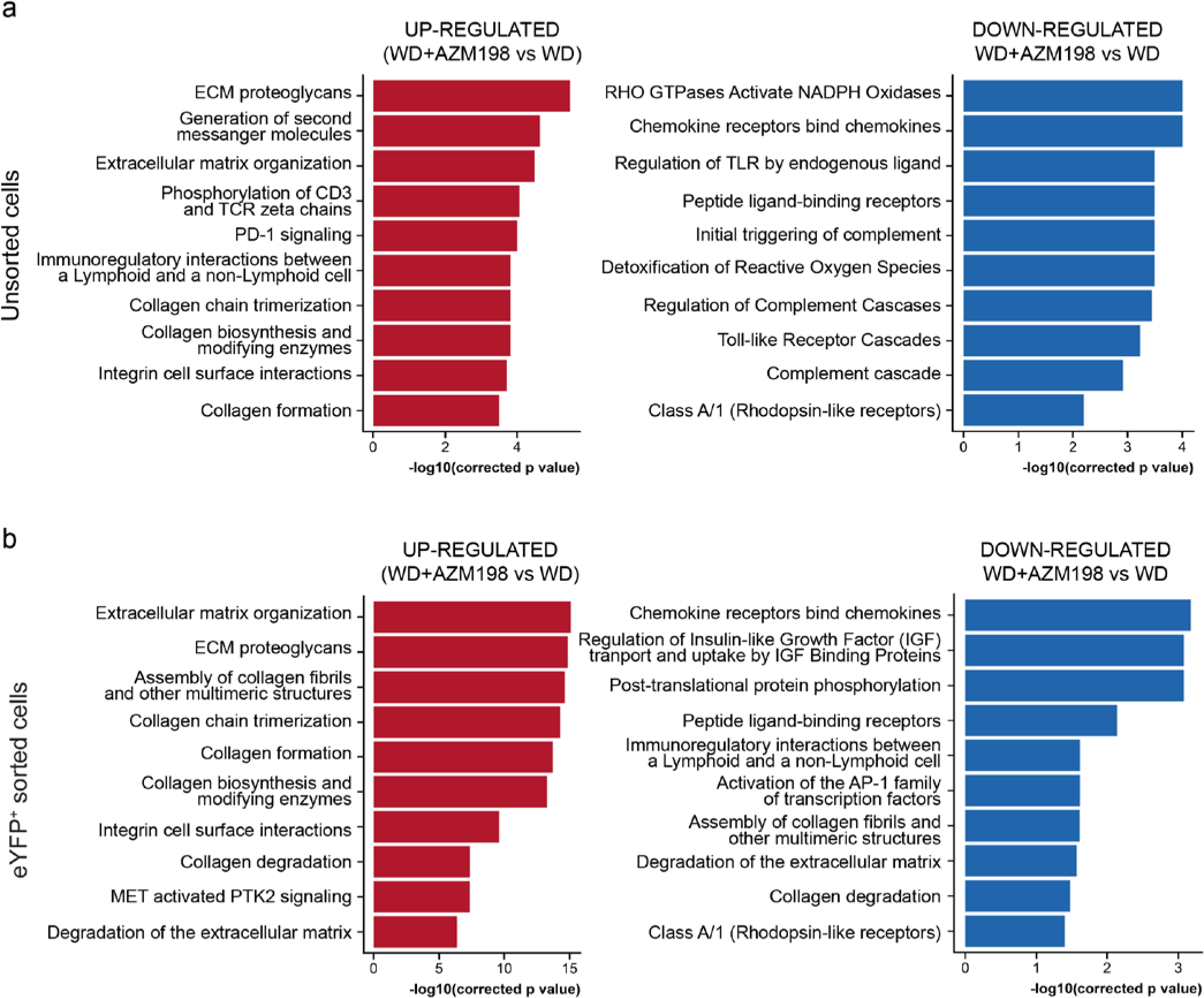
Reactome Pathway analysis from SR-BI^ΔCT/ΔCT^/*Ldlr^−/−^* mice fed WD+AZM198 versus WD. Samples of BCA regions from three mice for each condition (WD with or without AZM198) were pooled into one group each and subsequently analyzed. After generation of single cells from the tissue, samples of either the entire population of single cells (unsorted) or eYFP positive cells (SMC cells, eYFP+ sorted cells) were subjected to scRNA-seq analysis **(a)** The top 10 up-regulated pathways (left, red) and down-regulated pathways (right, blue) in the sample of unsorted cells from the BCA lesion region of SR-BI^ΔCT/ΔCT^/*Ldlr^−/−^* mice fed WD+AZM198 compared to those fed WD. **(b)** The top 10 up-regulated pathways (left, red) and down-regulated pathways (right, blue) in he sample of eYFP^+^ sorted cells from the BCA lesion region of SR-BI^ΔCT/ΔCT^/*Ldlr^−/−^*mice fed WD+AZM198 versus those fed WD.

## Methods

### Animal Handling and Tissue Processing

Mice were generated and used in accordance with protocols reviewed and approved by the Beth Israel Deaconess Medical Center, the Massachusetts Institute of Technology, and the University of Virginia Animal Care and Use Committees. Male *Myh11*-*CreER^T2^*/ROSA26 STOP^FL/FL^-eYFP^+/+^ mice (designated *Myh11-CreER^T2^*/Rosa-eYFP) have been previously described^67^. SR-BI^ΔCT/ΔCT^ mice^19^ were crossed with *Ldlr* knockout mice from The Jackson Laboratory (Strain# 002207) to generate SR-BI^ΔCT/ΔCT^/*Ldlr^−/−^* mice which we deposited to The Jackson Laboratory (Strain# 033709). SR-BI^ΔCT/ΔCT^/*Ldlr^−/−^* mice were then crossed with *Myh11-CreER^T2^*/Rosa*-*eYFP mice to generate experimental mice that were used for this study. The full genotype of the mouse is *Myh11-CreER^T2^*-eYFP SR-BI^ΔCT/ΔCT^ *Ldlr^−/−^* (abbreviated as “SR-BI^ΔCT/ΔCT^/*Ldlr^−/−^*” throughout). Exclusively male mice were used in all experiments since the *Myh11-CreER^T2^* transgene is located on the Y chromosome^68^. Mice were fed a tamoxifen diet (250 mg of tamoxifen/Kg diet) (Envigo; Catalog# TD.130856) *ad libitum* starting at 6-8 weeks of age to activate *Myh11*-driven Cre recombinase which removes a *loxp* flanked stop cassette in the ROSA*26* locus in front of an enhanced yellow fluorescent protein (eYFP) reporter gene allowing the expression of eYFP (**Figure. 1a**). As Cre excision is permanent, the *Myh11-CreER^T2^*/Rosa-eYFP/SR-BI^ΔCT/ΔCT^/*Ldlr^−/−^* mouse model provides permanent eYFP lineage tagging of virtually all *Myh11+* SMC that exist at the time of tamoxifen as well as their progeny. Following tamoxifen feeding and one week of rest (to ensure tamoxifen clearance), mice were fed standard rodent Chow diet (Teklad; Catalog# 7912), an atherogenic Western diet (WD) (containing 21% fat, 0.2% cholesterol) (Envigo; Catalog# TD.88137) or WD formulated to contain the MPO inhibitor AZM198 (500 µmol/Kg) from week 9-35 (total 26 weeks). AZM198 dosage was adopted based on studies done by AstraZeneca (Sweden)^7^. A daily AZM198 dose of 500 µmol/kg in WD was calculated based on an average daily food consumption of approximately 3.7 grams per mouse. Based on studies of human MPO purified from HL60 cells, this concentration corresponds to >95% inhibition of MPO activity^7^.

Mice were fed one of the above diets starting at 9 weeks of age *ad libitum* until they reached a humane terminal endpoint or for up to 26 weeks (35 weeks of age). Humane terminal endpoint was identified as exhibiting one or more of the following symptoms: decreased activity, ruffled coat, hunched posture, labored breathing, unsteady gait, hind limb paralysis and head tilt/spinning. Mice were carefully monitored at least twice daily by trained vivarium caretakers and researchers to recognize signs of MI. If any of the above signs were noticed during monitoring, mice were examined by trained veterinary technicians. If signs of MI or stroke or pain/distress were not able be alleviated by a veterinary technician, and euthanasia was recommended, this was considered the humane terminal endpoint. The animal was immediately euthanized by CO_2_ asphyxiation and tissues and blood samples were collected for analysis. Blood was used for complete blood count and plasma isolation. Then, mice were gravity-perfused via the left ventricle of the heart with 5 mL of PBS, 10 mL of 4% paraformaldehyde (PFA) (EMS; Catalog# 15710) in PBS, and an additional 5 mL of PBS. Organs were harvested, weighed, and digitally photographed. Tissues were fixed for 24h in 4% PFA in PBS with gentle shaking followed by processing and paraffin embedding.

### Echocardiographic Measurements

Serial transthoracic echocardiography was performed at indicated time points using a Vevo 2100 ultrasound system equipped with an MS400 probe (VisualSonics, Fujifilm). Mice were anesthetized with isoflurane in medical air throughout the duration of the procedure: 3-5% for induction and 1-2% for maintenance. At this dose of isoflurane, the heart rates were maintained at approximately 450 - 550 bpm. Once anesthetized, mice were placed on a heated platform that contained electrocardiogram (ECG) electrode pads. Chest fur was removed with a chemical hair remover, and ultrasound gel was applied. Both parasternal long axis and short axis images were acquired. Fractional shortening (FS) was measured from M-mode images of the short axis view. Analysis was performed using Vevo LAB software (VisualSonics, Fujifilm).

### Blood Analysis

For complete blood count (CBC), blood was collected in K_2_EDTA microtainer blood collection tubes (BD; Catalog# 363706). For plasma preparation, blood was collected in K_2_EDTA vacutainer tube (BD; Catalog# BD367835) and centrifuged. Plasma was stored at −80°C until analyzed for lipid profile (total cholesterol and triglycerides), multiplex cytokine analysis and for assaying liver toxicity enzymes – alanine transaminase (ALT) and aspartate transaminase (AST). Lipid profile, ALT and AST were analyzed by the University of Virginia Clinical Pathology Laboratory. MPO activity, multiplex cytokine analysis and AZM198 concentration in plasma were analyzed by AstraZeneca (Sweden) as described below.

### MPO Activity Assay

A high-binding, 96-well white plate (Lumitrac, PS flat-bottom; Greiner Bio-One; Catalog# 655074) was coated with 50 μL mouse anti-MPO capture antibodies (Abcam; Catalog# 16886; 1:200 dilution) and incubated overnight at 4°C. The plate was then washed three times with 350 µl PBS-Tween (PBS-T) and blocked with 1% BSA in PBS for 1 hour at 4°C. 50 μL plasma (diluted 1:3 in PBS) were added to each well followed by incubation for 2 hours at 4°C, at which time the plate was washed three times with 350 µl PBS-T. 50 μL Amplex Red Assay Buffer [50 mM NaPO_4_ pH 7.4, 140 mM NaCl, 10 mM Na_2_NO_2_, 40 μM Amplex Red (Invitrogen; Catalog# A12222), 10 μM H_2_O_2_] were added, and the plate was incubated in the dark for 25 minutes at room temperature with shaking and then fluorescence was measured using a SpectraMax Paradigm plate reader at excitation and emission wavelengths of 535 nm and 595 nm, respectively. Activity was calculated based on a standard curve determined using purified recombinant mouse myeloperoxidase protein (R&D Systems; Catalog# 3667-MP).

### Multiplex Cytokine Analysis

The multiwell assay plate, reagents, diluents, samples, and controls were prepared and used to detect the cytokine expression according to the manufacturer’s protocol (V-PLEX Plus Mouse Cytokine 19-Plex; Meso Scale Discovery, Rockville, MD, USA, Catalog# K15048G and K15245G). Briefly, the samples were prediluted 2.5x in kit sample diluent 41, and the standards were serially diluted to generate the standard curve. The Meso Scale Discovery (MSD) plate was washed three times with 200 μL wash buffer. The samples, standards, and quality controls (50 μL each) were transferred to the MSD plate and incubated for 2 hours at room temperature with gentle shaking. Afterward, the washing step was repeated. The detection antibody (25 μL) was added to plate and incubated for 2 hours at room temperature with gentle shaking. The washing step was repeated. 150 μL reading buffer were added and the values were recorded using MSD Sector Imager S600. Assays were performed using the Beckman Coulter robotic assay system (SCARA System) that incorporated a Biomek NX Span-8 and Biomek FX double 96-head pipetting robots, Cytomat2 shake incubator, Biotek EL405 washer, and MSD S600 reader.

### Plasma AZM198 Analysis

Plasma samples (20 µl each) were precipitated with 150 µl acetonitrile and vortexed, followed by a 20 min centrifugation at 3220xg at 4°C. Supernatants were transferred to a fresh 96 deep-well plate (Nunc 1mL; ThermoFisher Scientific; Catalog# 260252) and diluted 1:1 with water prior to LC-MS analysis. Matrix-matched calibration samples and blanks were prepared as were the study samples. Samples were analyzed using reversed-phase high-pressure liquid chromatography with rapid gradient elution. Compounds were detected using a Waters Xevo TQ-S triple quadrupole mass spectrometer (Waters Corporation, Milford, MA, USA). Chromatographic separation was performed using an ACQUITY BEH C18 1.7 µM, 2.1×50 mm column with the column temperature set to 40°C. The flow rate was maintained at 0.7 ml/min using a mobile phase of 2% acetonitrile and 0.2% formic acid in water (A) and 0.2% formic acid in acetonitrile (B). The elution gradient was ramped from 4% B to 95% B from 0 to 1.5 min, maintained at 95% B until 2.3 min and finally conditioned from 2.4 to 2.7 min. Mass spectrometry was performed using positive electrospray ionization and multiple reaction monitoring of transition 321.07 > 136.13 m/z.

### Histology

PFA-fixed paraffin-embedded tissues were sectioned to a thickness of 10 μm. For assessments of MI, the whole heart was sectioned from the apex to the aortic sinus. Following de-paraffinization and rehydration, every 5^th^ section (200 μm apart) was stained with Masson’s trichrome to identify myocardial fibrosis (indicative of MI), which stains fibrotic tissue blue and healthy myocardium red. Locations of the myocardial fibrosis were recorded. Fibrosis was quantified by ImageJ software (USA National Institute of Health) as number of blue pixels/sum of blue and red pixels. Myocardial fibrosis was expressed as total a percentage of total LV and RV area.

Coronary atherosclerosis was analyzed in Masson’s trichrome stained sections of the heart at the aortic sinus. Coronary lesion sizes and percent occlusion were quantified by ImageJ software as was done previously for BCA lesions^32,69,70^. To assess carotid artery and BCA morphometry, Modified Russell-Movat pentachrome staining was performed as previously described^32,69,70^. Plaque rupture in coronary artery, carotid artery and BCA was assessed both gross morphologically at the time of collection, and confirmed by Ter-119 staining of PFA-fixed paraffin embedded sections as previously described^32,46^. Coronary lesions were further assessed by immunofluorescence staining as described below.

For assessments of ischemic stroke, the fixed paraffin-embedded brains were sectioned (10 µm) from olfactory bulb to cerebellum. Following de-paraffinization and rehydration, standard hematoxylin and eosin (H&E) staining was carried out to determine the damaged area of the brain. The locations of the ischemic stroke in the brains were recorded. The brain damage due to ischemic stroke was further assessed by immunofluorescence staining using NeuN, Iba1 and GFAP antibodies as described below.

For liver histology, PFA-fixed paraffin-embedded tissues were deparaffinized and rehydrated. Liver morphology was assessed by H&E staining. Fibrosis in liver was assessed by Picro Sirius Red/Fast green (PSR) staining. For assessment of lipid deposition in the livers, Oil Red O staining was carried out. Briefly, PFA-fixed liver tissues were placed in 30% sucrose for 24 hours followed by embedding in OCT compound (Tissue-Tek; Catalog# 4583). OCT-embedded tissues were sectioned to a thickness of 10 μm. PSR and Oil Red O staining were carried out at the Histology Core at Cardiovascular Research Center, University of Virginia.

### Immunofluorescence Staining

For Immunofluorescence staining, sections were first deparaffinized in xylene and rehydrated in ethanol. Antigen retrieval was performed as per manufacturer’s instructions (Vector Lab; Catalog# H-3300). Sections were blocked in PBS containing fish skin gelatin (6 g/L) and 10% horse serum for 1 hour at room temperature. Tissues were then incubated with the following primary antibodies: GFP (Abcam; Catalog# ab6673; 1:100 dilution), MPO (Abcam; Catalog# ab208670; 1:200 dilution), FITC-conjugated ACTA2 (Sigma; Catalog# F3777 clone 1A4 1:500 dilution), NeuN (Abcam; Catalog# ab177487; 1:500 dilution), Iba1 (Fujifilm Wako; Catalog# 013-27691; 1:500 dilution), GFAP (Abcam; Catalog# ab7260; 1:500 dilution) or their isotype IgG as control at 4°C. Slides were incubated with DAPI (Invitrogen; Catalog# D21490, 5mg/mL; 1:100 dilution) and one of the following secondary antibodies: Donkey anti-rabbit Alexa Fluor 488 (Invitrogen; Catalog# A21206; 1:250 dilution), Donkey anti-goat Alexa Fluor 647 (Invitrogen; Catalog# A21447; 1:250 dilution), Donkey anti-rabbit Alexa Fluor 546 (Invitrogen; Catalog# A10040; 1:250 dilution) for 1 hour at room temperature. Following washing, slides were mounted with coverslips and Prolong Gold Antifade Mountant (Invitrogen; Catalog# P36930). Sections were imaged using a Zeiss LSM880 confocal microscope at either 20x or 40x magnification and 0.6x zoom to acquire a series of z-stack images at 1 μm intervals. Acquisition settings were determined using the IgG isotype control and were kept constant across images acquired. Maximum intensity projection was used to generate the representative images.

### Cell Processing and Data Analysis for scRNAseq

#### Cell processing

Four samples (unsorted and eYFP^+^ sorted cells from pooled BCA lesion area) from three WD-fed or three WD+AZM198-fed SR-BI^ΔCT/ΔCT^/*Ldlr^−/−^*mice were prepared as previously described and briefly summarized below^31,32,69^. Mice were euthanized by CO_2_ and perfused with 20mL of PBS + 1μg/mL Actinomycin-D. BCA lesion regions (as shown in Extended Data Figure 7a) were excised and placed into FACS buffer (1% BSA in PBS) on ice until all samples were collected. Tissues were chopped with scissors and digested in a cocktail containing 4 units/mL Liberase^TM^ (Roche; Catalog# 355374) and 0.744 units/mL Elastase (Worthington Bio. Corp; Catalog# LS002279) in RPMI + 1μg/mL Actinomycin-D for 60 min at 37°C. Individual samples were pooled (3 WD-fed mice pooled and 3WD+AZM198-fed mice pooled). A subset of pooled cells was taken for the “unsorted” sample and the remaining pooled cells were used for the “eYFP^+^” sample which were sorted using a BD Influx™ Cell Sorter. Live cells from “unsorted” and “eYFP^+^ sorted” samples were counted and submitted directly for scRNAseq library preparation.

#### Sequencing, read alignments, and quality control

All Libraries were prepared using the Chromium Next GEM Single Cell 3’ Kit v3.1 (v3 Chemistry) at the Genome Analysis and Technology Core at University of Virginia, in accordance with the manufacturer’s instructions. 400M reads/cell and ∼2000 cells per sample were targeted. Sequencing was performed on the NovaSeq 6000 sequencing system (Illumina) at Novogene, Sacramento, California, with system specifications of paired-end reads at 150bp length (PE150) and a total data output of 468 Gb.

FASTQ files were aligned using the cellranger count pipeline (version 6.0.1) to a custom mouse reference genome, mm10 (GENCODE vM23/Ensembl 98). This custom reference was established with cellranger mkref (version 6.0.1) by incorporating the FASTA and GTF files of the reporter gene (eYFP) into the mouse genome.

Data analysis was conducted in R (version 4.1.1) utilizing Seurat 4.3.01. WD groups (unsorted and sorted samples) exhibited a median of 11,949 and 15,300 UMIs per cell and 2,877 and 3,620 genes per cell, respectively. In contrast, the WD+AZM198 groups displayed medians of 6,922 and 18,919 UMIs per cell and 1,967 and 3,838 genes per cell, respectively. Uniform filtering parameters were applied to all samples. Cells expressing genes within the range of 420– 5800, denoting the 5th to 95th percentile, were considered. Cells with over 15% of reads originating from mitochondrial genes and over 5% from hemoglobin genes were categorized as poor quality and excluded.

#### Clustering, Annotation and Pathway Analysis

The gene counts were normalized using SCTransform^71^. To compromise between accuracy and computational needs of the dataset, two metrics were applied to select number of principal components (PCs), which were: 1) PCs that cumulatively contribute 90% of the standard deviation and 2) the point where the percent change in variation between the consecutive PCs is more than 0.05%. Based on analysis, the PC range spanned from 26 to 41, with 28 being chosen for clustering. To cluster and visualize all cells, a Louvain algorithm was used. ChooseR^72^ tool was used to determine the silhouette scores and identified range of resolutions from 0.4 to 0.8 as optimal for this dataset, and 0.6 was selected for further analysis. Assigning clusters identity was performed based on the canonical markers from literature-curated and statistically ranked genes. Differential expression analysis was performed using Model-based Analysis of Single-cell Transcriptomics (MAST^73^), and pathway enrichment analysis was based on REACTOME pathway database (ReactomePA version 1.44.0). Significantly enriched pathways were identified using a 5% false discovery rate cutoff, and the Benjamini–Hochberg method was used to adjust P values and their enrichment significance was quantified using −log10 of P value adjusted.

### Data Availability

The data and code that support the findings of this study will be made publicly available prior to publication.

### Statistical Analysis

Statistics were performed using GraphPad Prism 10.0.2 (GraphPad Software). Data are presented as mean ± standard error of the mean (s.e.m). Normality of data distribution was evaluated by the Shapiro-Wilk test and the *F* test was performed to check homogeneity of variance. For comparison of two groups of continuous variables with normal distribution and equal variance, two-tailed unpaired Student’s *t*-test (with additional Welch’s correction when variance was unequal) or for non-normal distribution, two-tailed nonparametric Mann-Whitney *U* test was performed. For comparison of three or more groups of continuous variables with normal distribution and equal variances, two-way analysis of variance (ANOVA) was performed followed by the Sidak’s method of multiple pairwise comparisons. Echocardiography results were evaluated by mixed-effects analysis followed by the Sidak’s method of multiple pairwise comparisons. A significance threshold of *P* ≤ 0.05 was used for all tests. The number of mice used for each analysis is indicated in the figure legends.

## Acknowledgements

The authors acknowledge Dr. Liam Rasch, Melissa Bevard, and Moon Snyder for their histology expertise, Rupa Tripathi for preparation of reagents required for enzymatic dissociation of cells for scRNA-sequencing, and Sultan Samim for assistance in monitoring mice for signs of decline. The authors also acknowledge the UVA Advanced Microscopy Core (RRID:SCR_018736) for training and use of the LSM880 confocal microscope and Leica Thunder Imager, the University of Virginia Flow Cytometry Core (RRID:SCR_017829) for flow sorting assistance, and the Genome Analysis and Technology Core (RRID:SCR_018883) for their roles in generating data in this manuscript. This work was partly funded by a research collaboration with AstraZeneca in addition to support from grant and HL 127174 to MK and OK, NIH R01 grants HL 156849, HL 155165, HL 141425 to GKO.

## Author Contributions

SS and RAD conceptualized and performed the bulk of the experiments, validation, data collection, analysis, and interpretation, and contributed significantly to development of methodology, manuscript writing, and editing. HD performed longitudinal echo and analysis of the associated data. MAE provided methodological help and analysis of brain tissue for stroke. SK and LSS contributed to scRNA-seq design, validation and collection of samples. AS contributed to the scRNA-seq data processing and analysis. VS, SB, GA contributed to data interpretation and manuscript editing. KW provided conceptual help in designing studies and played a role in data interpretation and manuscript preparation. OK and MK generated and performed preliminary characterization of the SR-B1^ΔCT/ΔCT^/Ldlr^−/−^ mice and played a role in data interpretation and manuscript editing. SH, NB, BK and MP supplied the AZM198 and contributed to data collection, analysis and interpretation. GKO supervised the entire project including assisting with conceptualization, experimental design, data interpretation, funding, and manuscript writing and editing. All authors viewed and approved the final version of the manuscript prior to submission. The authors declare the following competing interests: SH, NB, BK and MP are current or former employees of AstraZeneca (and own AZ shares). The MPO inhibitor used in this study (AZM198) was developed by AstraZeneca. AZ has an MPO inhibitor program in clinical development. GKO received partial funding for this project through a research alliance between AstraZeneca and the University of Virginia Robert M. Berne Cardiovascular Research Center. All other authors declare no competing interests.

Correspondence and requests for materials should be addressed to Dr. Gary K. Owens at

